# Mining for a New Class of Fungal Natural Products: The Evolution, Diversity, and Distribution of Isocyanide Synthase Biosynthetic Gene Clusters

**DOI:** 10.1101/2023.04.17.537281

**Authors:** Grant R. Nickles, Brandon Oestereicher, Nancy P. Keller, Milton T. Drott

## Abstract

The products of non-canonical isocyanide synthase (ICS) biosynthetic gene clusters (BGCs) have notable bioactivities that mediate pathogenesis, microbial competition, and metal-homeostasis through metal-associated chemistry. We sought to enable research into this class of compounds by characterizing the biosynthetic potential and evolutionary history of these BGCs across the Fungal Kingdom. We developed the first genome-mining pipeline to identify ICS BGCs, locating 3,800 ICS BGCs in 3,300 genomes. Genes in these clusters share promoter motifs and are maintained in contiguous groupings by natural selection. ICS BGCs are not evenly distributed across fungi, with evidence of gene-family expansions in several Ascomycete families. We show that the ICS *dit1*/*2* gene cluster family (GCF), which was thought to only exist in yeast, is present in ∼30% of all Ascomycetes, including many filamentous fungi. The evolutionary history of the *dit* GCF is marked by deep divergences and phylogenetic incompatibilities that raise questions about convergent evolution and suggest selection or horizontal gene transfers have shaped the evolution of this cluster in some yeast and dimorphic fungi. Our results create a roadmap for future research into ICS BGCs. We developed a website (www.isocyanides.fungi.wisc.edu) that facilitates the exploration, filtering, and downloading of all identified fungal ICS BGCs and GCFs.

## INTRODUCTION

Fungal specialized metabolites (SMs; also called secondary metabolites or natural products) are essential sources of antimicrobial (e.g., penicillin, griseofulvin) and therapeutic (e.g., cyclosporine, mycophenolate) compounds (1–3). In nature, SMs enable niche adaptation by conferring protection from abiotic and biotic stressors, as well as aiding in nutrient acquisition and competitive interactions (4). The bioactive properties of these molecules have propelled the research community to identify strategies to find new fungal SMs. Current genome-mining algorithms that predict SM-producing biosynthetic gene clusters (BGCs) in fungal genomes utilize the contiguous physical arrangement of genes to enable BGC predictions (5–7). This approach has successfully revealed millions of putative BGCs, many of which remain uncharacterized (8, 9). However, these algorithms rely on prior knowledge of chemical-class defining "backbone" synthases/synthetases (e.g., nonribosomal synthetases, polyketide synthases, terpene synthases/cyclases), biasing results towards well-characterized classes of BGCs. This approach cannot find BGC architectures with uncharacterized backbone genes (i.e., non-canonical). While there have been several recent efforts to predict non-canonical BGC architectures (10), the uncharacterized nature of non-canonical BGCs has made them challenging to incorporate into genome-mining software, obfuscating our broader understanding of their prevalence and function. Our inability to mine “unknown” BGCs (11) remains one of the greatest challenges hindering the discovery of novel compounds with bioactive properties (12).

Isocyanides (also called isonitriles) are a chemical class of SM produced by bacteria and fungi. These compounds are produced by non-canonical BGCs that are not detected by current genome-mining software (13). They are characterized by the presence of the highly reactive isocyanide functional group (R≡N^+^-C^-^), which is formed by the conversion of the amino group on select amino acids. Isocyanide SMs can participate in unique chemical reactions (14, 15) and possess potent antifungal, antibacterial, antitumor, and antiprotozoal bioactivity (16–18). The isocyanide functional group is renowned for its ability to chelate to metals (19, 20), and compounds with this group can preferentially bind to specific transition metals and metalloproteins (14, 17, 21).

The first isocyanide backbone enzyme, isocyanide synthase (ICS), was discovered by Brady and Clardy (2005) while investigating the metabolome of an unculturable soil bacterium. Since this initial finding, a small number of ICSs have been shown to serve as backbone synthases in bacterial and fungal BGCs (21, 23–25). The only three characterized fungal ICS BGCS are: (i) *xan* (17), (ii) *crm* (14), and (iii) *dit* (26–28). While the *dit1* (ICS) *dit2* (p450) cluster produces dityrosine, a core component in the cell wall of yeast ascospores that has no known metal association (26–28), the *xan* and *crm* BGCs are both strongly up-regulated under copper starvation (29). Despite the mounting evidence that ICS BGCs are a rich source of bioactive microbial natural products, many of which may have metal-associated properties (15, 17, 21), their exclusion from existing genome-mining software has limited research on them relative to other canonical classes of SMs. Lim et al. (2018) reported a diversification of ICS proteins across both the fungal and bacterial kingdoms. However, the chemical diversity of compounds that result from these synthases depends on the extent to which ICSs function as backbones in BGCs, as opposed to existing as standalone genes (30), and has yet to be fully elucidated.

In the present study, we addressed the hypothesis that ICS genes are core members in diverse fungal BGCs. We present the first genome-mining pipeline to identify eukaryotic ICS BGCs. We discovered 3,800 ICS BGCs in 3,300 fungal genomes, making ICSs the fifth-largest class of SM compared to canonical classes found by antiSMASH (6). In exploring this dataset, we focused on two unanswered questions: (i) What is the prevalence of ICS BGCs across different taxonomic groups? And (ii) what are the structural variations and commonalities among ICS BGCs in fungi (e.g., size, protein domain content)? Finally, we conducted additional analysis to examine the distinctive genomic and evolutionary signatures of the *dit1*/*2* gene cluster family (GCF) that was previously only described in yeast (referred to as the *dit* superfamily in this study).

## MATERIAL AND METHODS

### Dataset and genomic-database annotation

All publicly available annotated fungal genomes were downloaded from the NCBI database on 09/09/2021 using the NCBI’s Dataset tool, version 11.32.1. We generated protein domain predictions using HMMER v3.1b2 (e-value <= 1e-4) (31) with the Pfam database v34 (32). ICS proteins were identified based on the presence of the ICS-specific domain PF05141.1 (named ‘DIT1_PvcA’ in the Pfam database in reference to the *dit* cluster) (29). All canonical BGCs were predicted using the default settings within the fungal version of antiSMASH v5 (6). See ‘Step 1’ in the Reproducible Script for details on the code and specific parameters. See Supplemental Table 1 for information on the taxonomy and metadata of each genome.

### Locating putative ICS BGCs from regulatory-motif conservation

Genes within a BGC often share cluster-specific promoter motifs (i.e., genetic regulatory elements) that allow for the co-regulation of a cluster by a single transcription factor (33). To generate our BGC predictions, we examined the genes surrounding ICSs for shared promoter motifs using the eukaryotic BGC-detection algorithm, CASSIS v1 (34). All CASSIS predictions that contained one or more neighboring genes next to the ICS gene were considered BGCs and included in the subsequent analysis in accordance with past definitions of BGCs (7). Scripts detailing the implementation of CASSIS can be found in ‘Step 2’ of the Reproducible Script.

### Converting BGC predictions into standard file formats

Raw CASSIS predictions were run through a custom Python pipeline that converted each prediction into a GenBank (gbk), FASTA (fna), and General Feature Format (gff) file, enabling us to leverage existing BGC-processing software such as BiG-SCAPE (35), and cblaster (36). The raw gbk, fna, and gff files for all 3,800 ICS BGCs can be downloaded from the GitHub repository: https://github.com/gnick18/Fungal_ICSBGCs. Coding regions (CDSes) encoding SM-related biosynthetic protein domains (6) were flagged as ‘biosynthetic’ because BiG-SCAPE places more emphasis on these proteins when determining the relatedness of BGCs. The scripts used in this step can be found in ‘Step 3’ of the Reproducible Script. Refer to Supplemental Tables 2-4 for detailed summaries of the number of ICS BGCs in each species/genome and the protein domains present in each BGC.

### Grouping the ICS BGCs into GCFs

BGCs that belong to the same GCF are thought to produce identical or closely related natural products based on gene content and protein identity thresholds. ICS BGC gbk files were clustered into GCFs using BiG-SCAPE v1.0.1 (‘Step 4’ in Reproducible Script) (35). We created a modified ‘anchor file’ to ensure BiG-SCAPE treated each ICS gene as a chemical-class defining synthase (i.e., a backbone gene). Parameters for ICS GCF classification were tested using 13 different cutoffs ranging from 0.2 to 0.8 in increments of 0.05. Cutoff values above 0.45 were found to be too relaxed, leading to the merging of major ICS GCFs that were separate at lower cutoffs. Conversely, cutoffs below 0.25 were too strict, forming only small GCFs comprising BGCs from closely related species. A value of 0.3 (which is also the default value) was confirmed as the optimal cutoff. A network visualization of the BiG-SCAPE output was created with Cytoscape v3.9.1 (37) and can be found in Supplementary Figure S1. For more information on the dereplication pipeline run on the BiG-SCAPE output, refer to the Supplementary Methods.

### Identifying core genes in highly conserved ICS GCFs

CASSIS assumes that all the genes within a BGC share a cluster-specific promoter motif (34, 38). Although promoter-based BGC mining is an effective method for identifying fungal BGCs (6, 38), this approach fails to locate BGCs with differentially regulated genes (39). Therefore, we employed a second GCF-verification approach that relies on the co-localization of genes independent of regulatory motifs. Briefly, we searched for co-localized orthologous genes in each GCF to identify genes conserved across evolutionary time. The set of highly conserved genes within a GCF is referred to as a GCF’s biosynthetic core and represents the subset of genes essential for the biosynthesis of a given SM pathway (23, 40). Accessory genes, which are defined by their variability within GCFs, are thought to modify the SM produced by the biosynthetic core (4). Therefore, GCFs with the same core genes often produce related families of SMs (35). Several GCFs were identified as sharing a common biosynthetic core with other GCFs suggesting that these GCFs were not entirely resolved (9). In these instances, we collapsed GCFs with identical cores into a composite GCF that we call a ’superfamily’ and refer to original GCFs as ’Clans’ to clarify their distinct accessory gene complements. It was not feasible to perform manual steps of this analysis on all BGC predictions, so we focused on clusters present in the 31 ICS GCFs that contained at least five distinct species (core regions are also more easily identified among species). A detailed explanation of this methodology can be found in the Supplemental Methods. The refined BGC predictions can be seen in Supplementary Table 7.

### Estimating the extent of linkage disequilibrium in GCFs

We validated GCFs by confirming that the co-occurrence of genes found in BGCs within a GCF (i.e., linkage disequilibrium) could not be explained by random chance (i.e., genetic drift) (10). Whole-genome levels of linkage disequilibrium (LD) for species represented in GCF were calculated around BUSCOs (e.g., ‘Benchmarking Single Copy Orthologs’) (v3.0.3) (41) present across all species (*n* = 287). Around each BUSCO, we identified orthologous genes using Orthofinder v2.5.2 (default settings) (41) and calculated the percent conservation of orthologous genes across all species in a GCF as a proxy for LD. An identical process was used to estimate LD at ICS loci. The distribution of LD estimates around each BUSCO served as a null distribution to determine the significance of LD acting on ICS loci. The significance of LD was calculated using a percentile rank for each GCF (SciPy v1.5.2) (42).

### Phylogenetic inference by maximum likelihood (ML)

All maximum likelihood gene and protein trees were made using the following pipeline. First, sequences were aligned using MAFFT v7.475 with the ‘-auto’ parameter (43). Next, the alignments were trimmed using trimAl v1.2 with either the ‘-gappyout’ or ‘-gt’ parameters (44). Lastly, we made phylogenomic trees from the trimmed alignments using IQTree v2.0.3 (45) and ran 1,000 ultrafast bootstrap replicates (46). The optimal model of sequence evolution was selected using ModelFinder (47). All tree visualizations were constructed using either iTOL v5 (48) ggtree v3.1.1 (49), or ETE 3 v3.1.2 (50). All trees created for this publication can be found in the GitHub repository, along with the specific parameters used in their construction.

### Reconstruction of the Fungal Kingdom’s species tree

We chose a coalescent-based approach to construct a kingdom-level species tree because of this method’s computational scalability (51), compatibility with incomplete lineage sorting (52) and ability to handle missing data (53). We identified a set of 287 BUSCOs that were conserved across all 3,300 genomes (lineage database: ‘fungi_odb9’) (41). Unrooted maximum likelihood trees were generated for each BUSCO as defined above and used as the input for ASTRAL v5.7.8 (54). A species map file was generated and passed into ASTRAL using the ‘-a’ parameter to force isolates within the same species to be monophyletic. We verified that our reconstructed tree closely aligns with Li et al.’s (2021) established fungal tree of life. The tree was rooted at *Rozella allomycis*, the only publicly available genome from the earliest diverging lineage of the Fungal Kingdom, Cryptomycota (55, 56). BUSCO gene trees and the final species tree are available for download from the publication’s GitHub repository.

### Constructing a phylogeny of ICS backbone enzyme sequences

The phylogenetic relationships between GCFs were inferred from ICS backbone protein sequences using two datasets: 1) the ICS-domain sequence alone and 2) the entire protein sequence. Because both alignments produced topologically identical phylogenies, only the complete ICS protein alignment is presented in subsequent analyses.

### Creating the phylogeny of p450 proteins to gain insight into the relatedness of Dit2 orthologs

We investigated evolutionary relationships between Dit2 and other fungal p450 proteins to identify additional Dit2 orthologs. Constructing a phylogeny from every species’ set of p450 proteins was computationally prohibitive, as a single Ascomycete genome can contain hundreds of p450 genes. A subset of ten species was selected to represent the total diversity of the *dit* cluster. The species included: *Aspergillus clavatus*, *Candida tropicalis*, *Scheffersomyces stipitis*, *Saccharomyces cerevisiae, Monosporascus cannonballus*, *Kluyveromyces lactis*, *Rhizodiscina lignyota*, *Ophiobolus disseminans*, *Zygotorulaspora mrakii*, and *Macrophomina phaseolina*. Candidate p450s were identified in these genomes based on the presence of p450 protein domains. The resulting dataset of 845 p450 protein sequences was constructed into a maximum likelihood phylogeny as defined above.

### Topological comparison of the Dit1/2 phylogeny and the reconstructed species tree

Tests of phylogenetic incongruence were used as they are considered the most reliable way to infer historical events from genomic and proteomic sequences (57, 58). Before creating a concatenated phylogeny from the Dit1 (ICS) and Dit2 (p450) protein sequences, we verified that phylogenies inferred from these proteins had similar topologies, suggesting that they had co-evolved. The individual protein trees were topologically aligned using the R packages ape v5.6-2 (59), phangorn v2.8.1 (60), and TreeDist v2.4.1 (61), and Robinson-Foulds distances were calculated (62, 63). The optimal model of sequence evolution was estimated independently for both genes using ModelFinder and found to be LG+I+G4 in both cases (47). After verifying the co-evolution of the Dit1 and Dit2 proteins, we concatenated these sequences using FASconCAT-G v.1.05.1 (64) and repeated the process, performing a topological comparison between the concatenated Dit1/2 tree and the species tree to determine if this GCF’s evolutionary history could be explained by vertical transmission.

## RESULTS AND DISCUSSION

### ICS BGCs are not evenly distributed across the Kingdom Fungi

Our ICS-BGC-detection pipeline (see Methods) identified 4,341 ICS genes within 3,300 genomes representing 1,329 unique species (Figure 1, Supplementary Table S1). Cluster-specific promoter motifs were found in genes surrounding 3,800 (87.5%) of the 4,341 ICS genes. After selecting a single representative for each species, the ratio of ICS proteins in BGCs remained very similar, with 1,190 BGCs found around 1,394 detected ICS proteins (85.4%). This data suggests that most ICSs function as backbone enzymes in fungal BGCs (Supplementary Tables S2-4).

**Figure 1.**
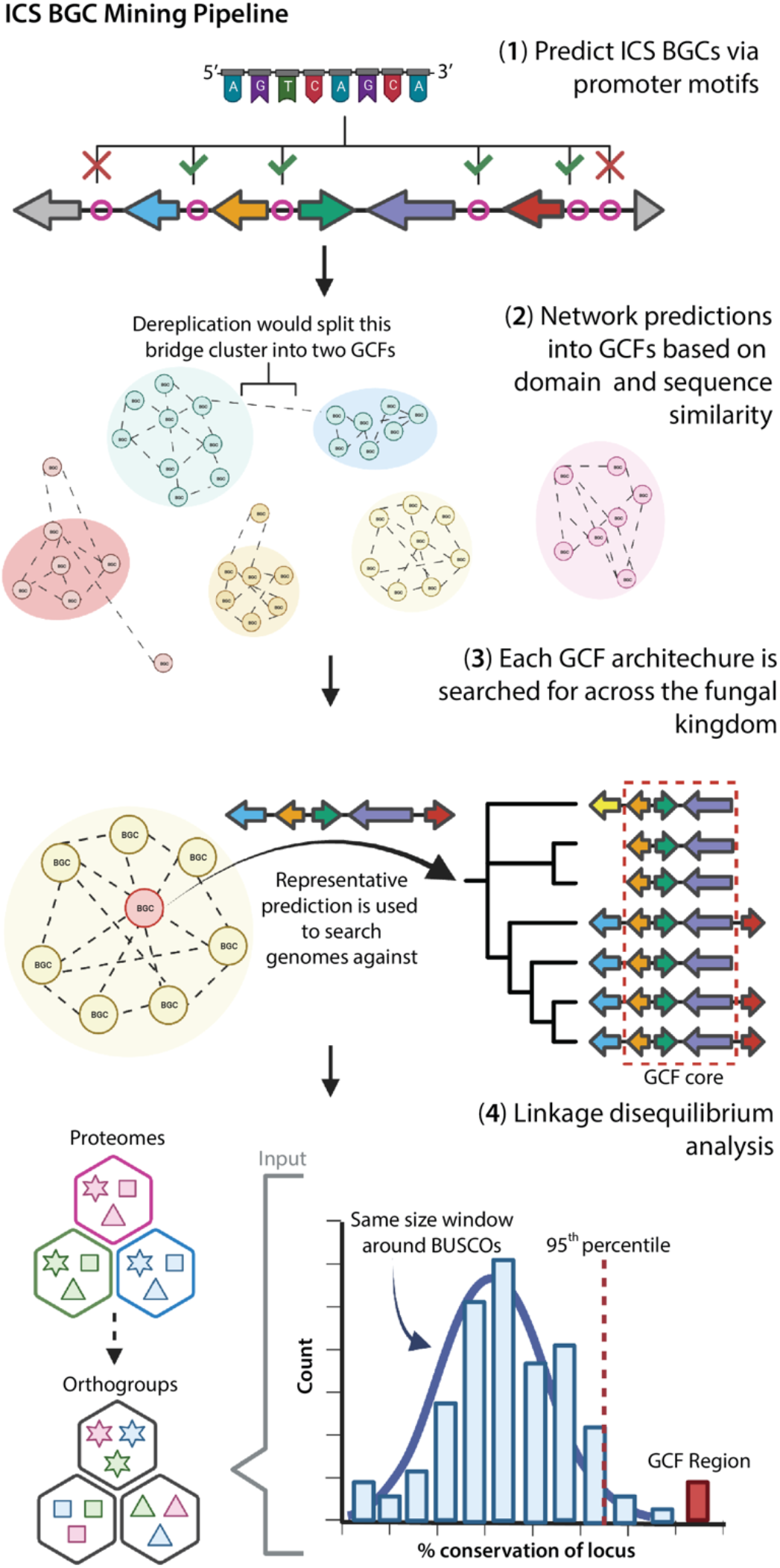
Full isocyanide synthase (ICS) biosynthetic gene cluster (BGC) genome-mining pipeline. First, clusters of co-regulated genes around ICSs are predicted. These BGCs are then grouped into gene cluster families (GCFs) based on sequence similarity and shared protein domains. To identify the core genes in GCFs, we searched for co-localized orthologous genes across every genome in the dataset. Finally, a subset of the GCFs was validated by confirming that the co-occurrence of genes found in the GCFs (i.e., conservation of genes in all GCF BGCs) exhibited significant linkage disequilibrium (LD) relative to whole-genome levels of LD (i.e., conservation of genes around BUSCOs). To determine if two genes were evolutionary related, all genes were first grouped into orthologous gene groups (i.e., orthogroups).

The genomes of species within the Ascomycete classes Sordariomycetes, Dothideomycetes, and Eurotiomycetes contained the most ICS BGCs. However, the genomes of some individual species outside of these groups were also found to contain several ICS clusters; for example, the genome of the saprobic Basidiomycete species *Tetrapyrgos nigripes* contained 6 ICS BGCs and the coral mushroom *Clavulina* sp.’s genome contained 5 ICS BGCs (Figure 2; Supplementary Table S2). A preliminary examination of the expansion of ICS BGCs across the phylogeny of Ascomycota identified 17 branches where sister lineages showed significant differences in the number of clusters (Supplementary Figure S2; Supplementary Methods), suggesting a gene-family expansion and/or contraction in these lineages. For example, genomes in one lineage of the genus *Colletotrichum* contained an average of 4 ICS BGCs, while a sister lineage had an average of 1 ICS BGC. Plant pathogenic taxa, such as *Penicillium*, *Bipolaris* (*Cochliobolus*), *Colletotrichum*, and *Fusarium* (65, 66), were also found to harbor the greatest number of ICS BGCs. Future research should focus on determining the ecological drivers of ICS gene cluster expansions and contractions in these lineages.

**Figure 2.**
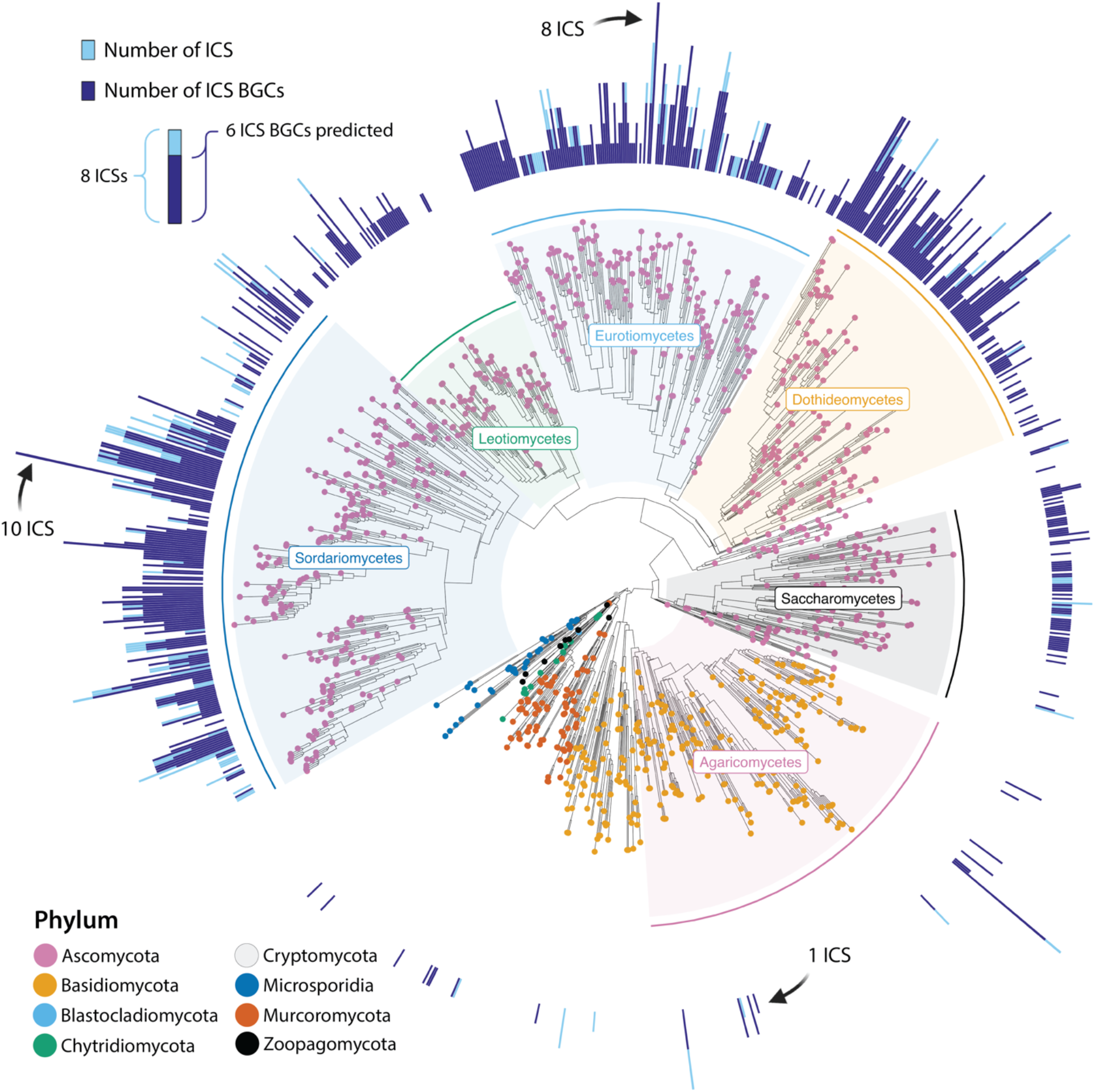
Distribution of isocyanide synthases (ICSs) and associated clusters (BGCs) across the fungal kingdom. The species tree was estimated using a multi-species coalescent model, and is rooted at the early-divergent subphylum, Cryptomycota. Tips are colored based on phylum, and major classes within the Ascomycota and Basidiomycota are labeled. The overlay bar chart on the outer ring depicts the number of ICS genes found (dark blue), and number of ICS BGCs (light blue). Fully dark blue bars denote that all ICS genes act as backbones in ICS BGCs, while fully light blue bars indicate that none of the predicted ICS genes were found in co-regulated gene clusters.

ICS BGCs accounted for 3% of the Fungal Kingdom’s biosynthetic potential, with an additional 84,657 canonical BGCs identified by antiSMASH (Supplementary Figure S3) (6). Proportionally, this makes ICSs the fifth-largest class of predictable fungal BGCs. While siderophores are the most prominent group of SMs that facilitate microbial interactions with metals (67, 68), we discovered three times more ICS BGCs than siderophore BGCs. Much like the metal-regulation of siderophore synthesis (69), both characterized isocyanide natural products in *Aspergillus fumigatus* were recently shown to be tightly regulated by external copper concentrations. The isocyanide, xanthocillin, synthesized from the *xan* BGC not only provides chalkophore-like properties for *A. fumigatus* but also inhibits growth of bacteria and other fungi by withholding copper from these microbes (17).

Similar metal-associations have been found with bacteria isocyanides (2, 21). A bacterial isocyanide, rhabduscin, decorates the cell surface of some gram-negative bacteria and enables infection of insects by inhibiting the activity of central immune system copper-dependent metalloenzymes (15). The prevalence of fungal ICS BGCs, along with evidence of their metal associations, raises questions about the potential importance of this class of SM in mediating metal-associated fungal ecology and pathogenicity, emphasizing a new opportunity to expand our understanding of how fungi interact with and utilize metals.

While most taxonomic families contained 0-2 ICS BGCs, the right tail of the ICS BGC distribution among taxa comprised 12 families with 4-7 ICS gene clusters (Supplementary Figure S4a-b). In this group, ICSs accounted for 7-15% of the species’ total SM BGCs (Supplementary Table S5). Overall, the number of ICS BGCs within a genome had a strong positive correlation with the number of BGCs (canonical & ICS, Pearson’s coefficient: 0.39; *p*: 2.09 x 10^-75^). Part of this association may reflect that larger genomes (as defined by gene counts) have more BGCs (Pearson’s coefficient: 0.30; *p:* 1.86 x 10^-26^) (Figure 3a). We also investigated the impact of genomic sequence quality on the number of BGCs and observed a strong positive correlation (Supplementary Figure S4c; Pearson’s coefficient: 0.22; *p*: 1.14 x 10^-14^). While several studies have posited ecological drivers for the number of clusters in genomes (10, 70), they have not counted for genome size. Our finding suggests that the number of BGCs in a genome may, to some degree, correlate with size of the genome or genomic sequence.

**Figure 3.**
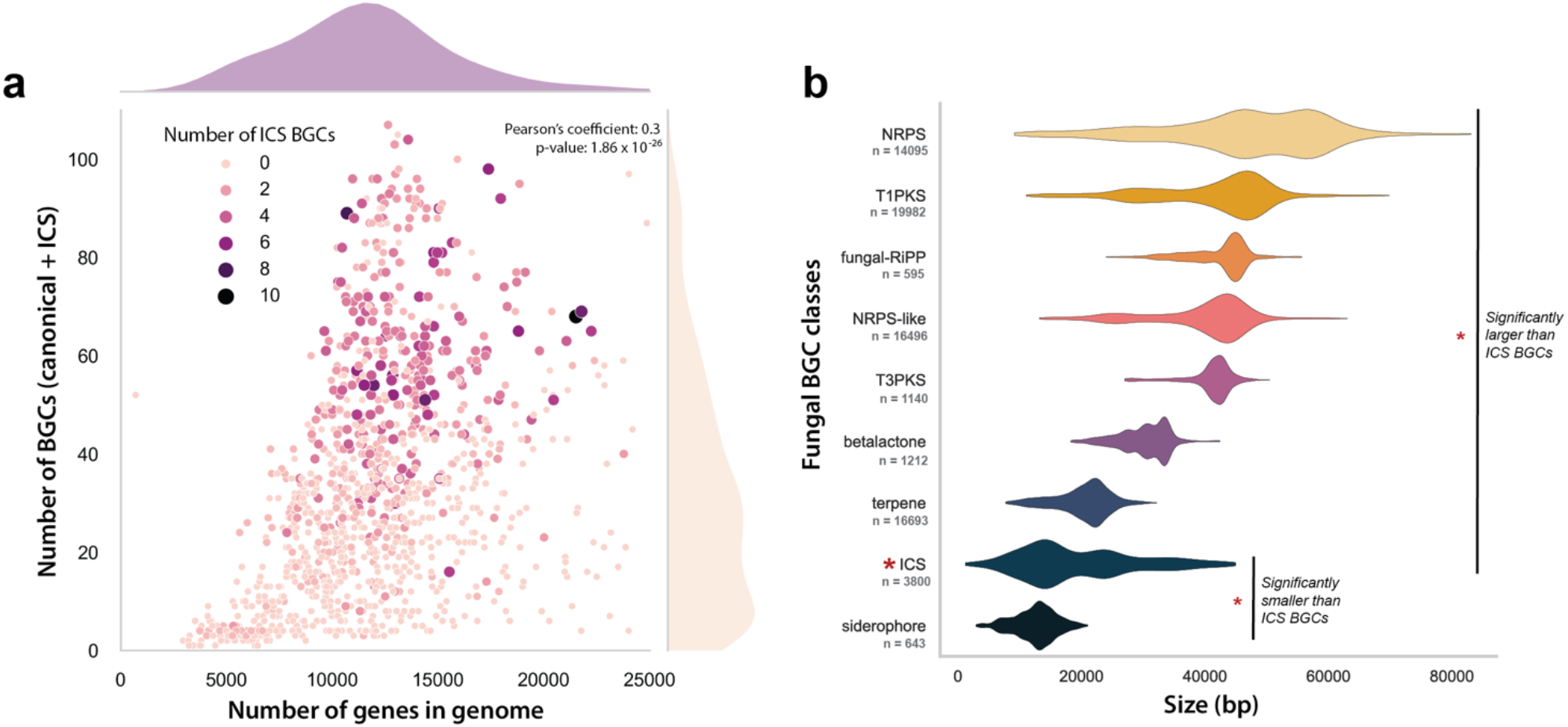
Exploring the relationships between isocyanide synthase (ICS) biosynthetic gene clusters (BGCs), canonical BGCs (predicted by antiSMASH), and genome sizes. (a) A scatterplot of the total number of BGCs (canonical + ICS) and the number of genes in a genome. Each point represents a genome and is colored and sized based on the number of ICS BGCs predicted within the genomes. Density plots are added to the margins of the scatterplot to show the distribution of the datapoints along the x and y axes. (b) The distribution of BGC sizes (in base pairs) for canonical BGC classes, and the ICSs elucidated in this study (*n* = BGCs). BGC classes that were significantly different in size (T-tests) from ICS BGCs are marked on the plot.

### The structural diversity of ICS proteins and ICS BGCs

All ICS enzymes in our dataset matched one of three protein domain variations summarized by Lim et al. (2018) (Figure 4b): monodomain ICSs (27), two domain ICS-dioxygenases (ICS-TauD) (17), and the multidomain ICS-nonribosomal peptide synthase-like (ICS-NRPS-like) variants (14). The relative abundance of each backbone-gene variant across species (with only a single genome representing each species) was approximately equal. ICS-TauD, ICS-NRPS-like, and ICS monodomain accounted for 36.2%, 32.8%, and 31% of the total ICS proteins, respectively. Outside of the backbones, 55.5% of clustered genes encoded protein domains associated with SM biosynthesis, transportation, or regulation, 22.7% encoded protein domains not typically associated with SM biosynthesis (e.g., sugar transporters, membrane proteins, RNA-related enzymes), and 21.8% lacked a predictable domain altogether.

**Figure 4.**
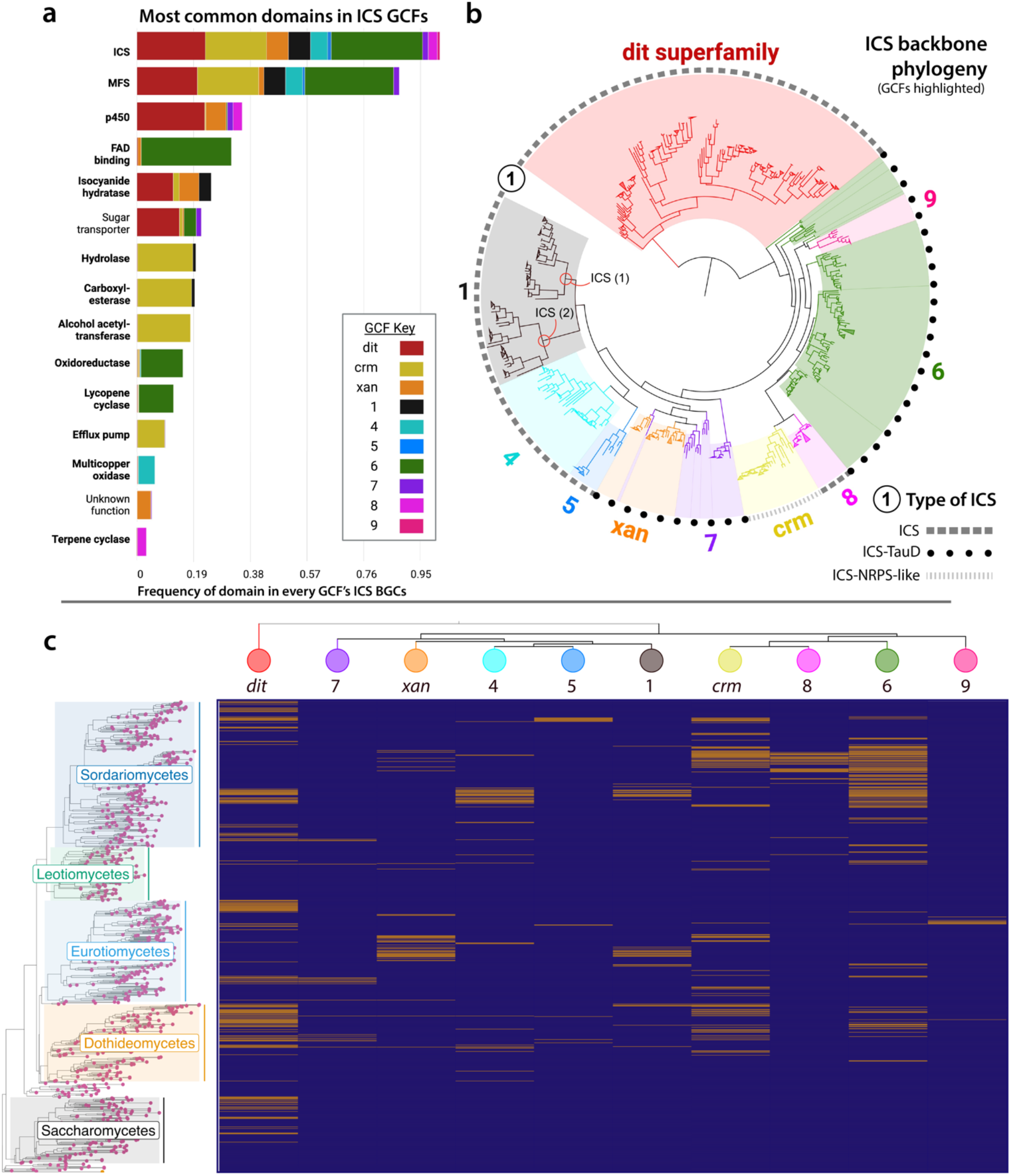
Architecture, evolution, and distribution of isocyanide synthase (ICS) gene cluster families (GCFs). (a) A subset of the most common protein domains in the ICS GCFs. Domains frequently found within ICS backbones variations (e.g., TauD, NRPS-like) were filtered out, except for the ICS domain (top row) which is present in 100% of the clusters. GCFs are labeled with colors (see key), and domains implicated in specialized metabolism biosynthesis, transportation, or regulation are bolded. An un-subset version of the plot can be found in Supplementary Figure S11. (b) Midpoint rooted maximum likelihood phylogeny of ICS proteins from the analyzed GCFs. The tree is colored and labeled with the same key found in section ‘a’. The outer black ring conveys the variation of the ICS enzyme based on its dashed pattern. (c) Presence (yellow) and absence (purple) heatmap of each GCF mapped onto a species tree of Ascomycota. Columns are sorted by the relatedness of each GCF’s ICS backbone using a collapsed ICS protein phylogeny.

ICS BGCs are significantly smaller than all other classes of SM BGCs except for siderophores. ICS BGCs have a mean length of 20,589 bp and contain an average of 8.42 genes per cluster (Figure 3b; Supplementary Figure S5). The small size of ICS BGCs may suggest that resulting compounds are subject to less chemical modification by tailoring enzymes than compounds resulting from other classes of BGC, although we cannot rule out that differences in BGC size between SM classes could result from biases in BGC-border prediction defined by genome-mining algorithms. The observation that ICS BGCs tend to be small is consistent with what has been found in characterized fungal ICS BGCs, with dityrosine production in *S. cerevisiae* requiring only two genes (27) and the *xan* BGC in *A. fumigatus* containing only five genes (17). However, small BGCs can generate outsized chemical diversity through trans-BGC interactions. We speculate that the highly reactive isocyanide backbone may be more likely to have such interactions. Indeed, the ICS product of the *crm* BGC can react with an ergot alkaloid precursor produced by another BGC, resulting in a novel class of alkaloids known as the fumivalines (14). Such reactions could dramatically increase the chemical diversity generated by ICS BGCs despite the relatively small size of these clusters. The nucleophilic, electrophilic, and metal-ligand properties of isocyanides (16) that have made them widely utilized in synthetic (71, 72) and medicinal chemistry (20, 73), suggests exciting prospects for exploring their potential biological activities.

### ICS biosynthetic diversity across taxonomic ranks

We dereplicated ICS BGCs into GCFs (Supplementary Figure S1) (8–10) to clarify the biosynthetic potential that these clusters represent. Of the 3,800 ICS BGCs, 92.4% could be categorized into GCFs (Supplementary Table S6), while the remaining clusters were found only in a single genome. Of the 200 GCFs we identified, 47.3% (*n* = 90) were detected in multiple species, while 52.7% (*n* = 110) were found only within isolates of the same species (Supplementary Figure S6; Supplementary Table S7). In rare instances, we observed multiple copies of the same GCF in individual genomes. While on average the ICS GCFs were highly conserved within species (98.6%), we located 1,963 instances where intraspecies isolates shared less than 50% of the same ICS GCFs. While intraspecific differences in SMs have historically been treated as random variation, recent work has shown that such differences can help to explain niche adaptation and macroevolutionary processes (74). More work is needed to clarify if intraspecific variations in ICS BGCs are associated with specific lineages within species. The vast majority of GCFs found across multiple species were unique to a single genus (70%). Although a small portion of the GCFs we identified (*n* = 22 or 11.5%) was found in two or more taxonomic families, these GCFs were highly conserved across the Fungal Kingdom and accounted for a disproportionate 40.7% (1547/3800) of all ICS BGC predictions.

### The distribution and evolution of ICS GCFs in fungi

Understanding the evolutionary history of fungal GCFs presents a challenge because different GCFs frequently lack a shared set of orthologous genes (75, 76). However, by definition, backbone genes (and or enzymes) are conserved across GCFs of a given biosynthetic class and can thus be used to infer the evolution of these clusters (77). We investigated the phylogenetic relationships of ICS backbone proteins from a subset of 10 ICS GCFs (see Methods), with the goal of determining whether the gene combinations that define BGCs within a GCF shared a single origin or whether they evolved convergently. Most GCFs could be mapped to monophyletic clades of backbone enzymes, indicating that these GCFs had a single common ancestor (Figure 4b). The ICS enzymes from GCFs 9 and *xan* were nested within GCFs 6 and 7, respectively. This suggests the ICS proteins in GCFs 9 and *xan* may have evolved from ICS within GCFs 6 and 7, respectively, sometime after the latter GCFs had already evolved. However, these BGCs do not share any genes beyond their ICSs. This pattern could be explained by the translocation of an ICS backbone gene from a parent GCF into a new genomic context. However, despite strong bootstrap support for the topology of these branches (Supplementary Figure S7), we cannot rule out that these relationships are not fully resolved in our phylogeny, as single-protein trees offer relatively little phylogenetic resolution. The two ICS backbone enzymes found within GCF 1 formed two distinct clades stemming from a single common ancestor (Figure 4b). A small clade of ICSs intercedes the two aforementioned clades in the full ICS phylogeny (see full ICS phylogeny in GitHub repository), however, none of the species represented in this clade have GCF 1. This observation suggests that the two enzymes in GCF 1 arose from an ancestral duplication event, which has been conserved over evolutionary time.

The three ICS protein-domain variations (see structural diversity section above) did not form monophyletic groupings, raising questions about the possibility of multiple emergences of the two variants with discontinuous distributions: the ICS-TauD and monodomain ICS variations (outer ring on Figure 4b). A parsimonious explanation is that an ICS-TauD ICS was the common ancestor of the ICSs found in GCFs 6, 7, and 9 and that multiple loss-of-function mutations led to the evolution of the monodomain ICS found in GCFs 1, 4, 5, and 8. However, we cannot rule out the possibility that the ICS-TauD could have independently emerged once in the ancestor of GCF 7 & *xan* and again in the ancestor of GCFs 6 & 9. It is also possible that the observed discontinuous patterns of ICS domain variations result from hemiplasy caused by incomplete lineage sorting, rather than gene loss or independent emergences (78). The *crm* GCF is the only example of an ICS-NRPS-like backbone gene (14). However, the GCF with an ICS that is the closest relative of *crm* (GCF 8) contains a monodomain ICS backbone that is co-localized with an NRPS gene containing a domain identical to that found in the *crm* ICS-NRPS-like proteins (Fig. S4). A parsimonious interpretation is that a mono domain ICS fused with a nearby NRPS in some lineages, but that these genes remained separate in an ancestral *Fusarium* species.

We observed three main patterns of distribution among GCFs. GCFs 5, 8, and 9 had narrow distributions and were primarily present in 1–2 closely related genera (Figure 4d). Similar distributional patterns of canonical GCFs are thought to reflect the conserved functionality of a GCF that may help define the ecological niche of closely related taxa (1, 79–81). GCFs *xan*, 1, 4, and 7 had discontinuous distributions (Figure 4d) but were more broadly spread across taxonomic groups. Three GCFs—*dit*, *crm*, and 6—were widely distributed across Ascomycota (Figure 4d). While these latter GCFs are not found in every lineage (Supplementary Figure S8), similar distributional patterns of the pseurotin gene cluster have recently been explained with vertical transmission – a pattern that emphasizes the potential importance of incomplete lineage sorting and gene loss even in SMs that often have strong ecological phenotypes (82). The latter three GCFs are widespread and account for 27.5% of all fungal ICS BGCs. For example, ICS BGCs from GCF 6 were present in the genomes of Eurotiomycete, Sordariomycete, Leotiomycete, Dothideomycete, and a single Basidiomycete species. Previous studies have suggested that BGCs involved in essential biological functions, such as growth and development, tend to be highly conserved and broadly distributed (76, 83). The distributional patterns we identify offer insights into the niche overlap of these lineages and raise questions about evolutionary forces maintaining these genes in specific lineages.

### Protein domains found within ICS GCFs

Canonical BGCs contain a core synthase and/or synthetase, as well as tailoring enzyme genes (4), and frequently include genes for transporters, protective enzymes, and transcription factors (70, 81). Available evidence suggests that ICS BGCs share many of the same genomic signatures, as all characterized bacterial and fungal ICS BGCs contain both a backbone synthase and tailoring enzymes (17, 21, 25, 26). However, the importance of specific protein domains in the function of ICS BGCs remains poorly understood. To identify protein domains associated with the evolution of ICS BGCs, we analyzed protein domains of all 3,800 ICS BGCs and within a subset of 10 ICS GCFs (see Methods; Figure 4).

Major facilitator superfamily (MFS) genes, which are typically associated with substrate transport (84), were the most common type of gene (after ICS backbones) in ICS BGCs, with approximately 60% of all ICS BGCs containing at least one MFS (Supplementary Table S8). The following three most common genes encoded cytochrome p450s (34.6%), zinc finger transcription factors (21.3%), and isocyanide hydratases (21.1%), the latter hypothesized to provide self-protection from isocyanide moieties (85). For a complete list of the most common protein domains found in all the ICS BGCs, refer to Supplementary Table S8.

We then looked at the biosynthetic core of 10 sampled GCFs. A GCF’s biosynthetic core is highly conserved and typically represents the minimum set of genes essential to produce a given SM pathway (23, 40). Therefore, we hypothesized that the ICS GCF biosynthetic cores would contain a higher proportion of genes associated with SM biosynthesis, transportation, or regulation than all the genes present within the ICS BGCs. Of the 57 core genes examined, 48 (84.2%) were found to be associated with SM production (Supplementary Figure S9b; Supplementary Table S9). This proportion is nearly 30% higher than that of biosynthetic genes detected in all ICS BGC genes (Supplementary Figure S9a). The most abundant genes in GCF cores were also the most common overall genes (encoding MFSs, p450s, and isocyanide hydratases). The functionality of other core genes was specific to individual GCFs. These genes encoded alpha/beta-hydrolases, alcohol acetyltransferases, oxidoreductases, efflux pumps, lycopene cyclases, multicopper oxidases, and terpene cyclases, among others, as shown in Figure 4a, Supplementary Figure S10, and Supplementary Figure S11. Overall, many of the commonly co-opted genes are similar to those found in canonical fungal BGCs.

Lastly, we checked for the presence of ICS BGCs that also contain a canonical synthetase and or synthase. BGCs containing two or more synthetases/synthases are not uncommon in the fungal kingdom (*n* = 86) but can contribute to the chemical complexity of SMs (*n* = 87). We focused our search on ICS BGCs containing NRPS, PKS, or terpene domain proteins (Table 1; Supplementary Methods). We found 12.8% (*n* = 685) of all ICS BGCs had genes encoding various NRPS domains. Most of these cases were explained by ICS-NRPS-like variants within GCF *crm* that commonly contain an ICS domain, an AMP binding domain, a phospho-pantethine (PP) loading domain, and a transferase domain (*n* = 14). A total of 95 ISC BGCs contained genes encoding both NRPS-like proteins and terpene synthases; the majority (89.5%) of these cases were found in the *Fusarium*-specific GCF 8 (Supplementary Table S10). A small number of BGCs (*n* = 12) also contained a putative PKS within the ICS BGC, primarily in the genomes of select Dothideomycetes and Sordariomycetes species. Although multiple backbone genes have been shown to interact to form a single SM (88), in other cases, BGCs with multiple backbone genes have been shown to encode distinct products (89). Whether or not these genes encode activities that contribute to ICS chemistry is yet to be determined.

**Table 1.** The proportion of each biosynthetic gene cluster (BGC) class across all fungal genomes. ICS BGCs are bolded. Multiple isolates within the same species were filtered to account for the overrepresentation of specific species within the dataset. * The siderophore BGC class is composed of both NRPS and PKS siderophore pathways (per antiSMASH classifications) *\*\** The ‘Other’ row encapsulates all fungal BGC classes smaller than fungal-RiPPs

| BGC Classes | Percent |
| --- | --- |
| Nonribosomal peptide synthase (NRPS) | 34.60% |
| Type 1 polyketide synthase (T1PKS) | 22.70% |
| Terpene | 22.40% |
| NRPS,T1PKS | 6.40% |
| **Isocyanide synthase (ICS)** | **3.00%** |
| Indole | 2.80% |
| Type 3 polyketide synthase (T3PKS) | 1.40% |
| Betalactone | 1.10% |
| Siderophore* | 0.90% |
| Fungal ribosomally synthesized, post-translationally modified peptide (fungal-RiPP) | 0.40% |
| **Other | 4.30% |

### Selection maintains ICS GCFs

We validated BGC predictions using the principles of linkage disequilibrium (LD) to test the hypothesis that the co-localization of genes in BGCs is most likely explained by selection, not genetic drift (see Methods) (10). Between species, LD can be impacted by recombination rates, genetic drift, sampling, and selection. However, we aimed to isolate the impact of selection by comparing LD estimates within BGCs to estimates made across the full genome (we assume that drift and recombination impact the whole genome). This approach found significant LD maintaining the structure of three known ICS GCFs: *dit*, *crm*, and *xan*. Based on these results, we focused our search on a small subset of 15 out of the 200 GCFs (refer to Methods section on estimating LD). This subset was necessary to capture the most taxonomically widespread ICS GCFs because high LD found within species and between closely related species offers little resolution to identify genes under selection. Two of the GCFs exhibited strong LD (*p* < 0.07) and 12 GCFs had significant LD (*p* < *0*.05) (Supplementary Figure S12). A single GCF primarily present in *Colletotrichum* species (GCF 26 in Supplementary Figure S12n) exhibited no significant LD patterns (*p*: 0.15). In this GCF, approximately 75% of BGC predictions included a core set of 7-10 SM-related genes (e.g., transcription factors, tailoring enzymes, transporters), while the remaining 25% contained only 1-3 genes. Upon manual inspection, we identified that the smaller BGC predictions were colocalized with orthologs found in the larger predictions. However, these genes did not share the same promoter motif as was found around the ICS backbone gene. These results suggest that either new genes have been recruited to the cluster in some lineages or, conversely, these genes are no longer coregulated with the ICS. Different regulatory patterns between species, like those we inferred here, have previously been associated with ICS gene clusters. The cluster-specific transcription that is part of and regulates the ICS *xan* cluster in *Aspergillus fumigatus* instead regulates the citrinin gene cluster in *Penicillium expansum* where the *xan* promoter motif has been lost (39).

The 14 validated GCFs account for nearly 50% of all fungal ICS BGCs (1865/3800). While we cannot entirely rule out other factors that influence LD, the significant co-occurrence of the same orthologous genes in these ICS GCFs relative to whole-genome levels of LD strongly suggests that selection is maintaining the structure of ICS GCFs. Similar evidence has also been used to infer the selection of BGCs by Gluck-Thaler et al. (2020). The physical clustering of genes into BGCs is often attributed to the stepwise process of SM biosynthesis. The backbone gene synthesizes a core chemical moiety that is then adorned by tailoring enzymes. Pathway intermediates may be passaged through cellular organelles (e.g., peroxisomes) by transporters and the final SM product may then be exported from the fungus through additional transporters. The loss of any gene in this process can disrupt SM synthesis, even leading to toxic impacts on the producing fungus (90). While we were unable to validate LD acting on GCFs with narrow taxonomic distributions or differentially evolved regulatory patterns, these results validate our approach to identifying ICS gene clusters and emphasize that these clusters are maintained by strong natural selection.

### The *dit* superfamily is widely distributed across Ascomycota

The *dit* cluster (27) is the archetypal ICS cluster for which the associated protein domain is named and defined in protein-domain-finding algorithms used here (29, 32). In yeast, this cluster is made up of two genes, *dit1* (an ICS) and *dit2* (a p450), that together synthesize dityrosine. Dityrosine is a structural component required for the normal formation of ascospore cell walls in *Saccharomyces cerevisiae*. This compound has only been reported in the yeasts *S. cerevisiae* (27) and *Candida albicans* (91). While a recent study raised questions about if this cluster encodes a true ICS (92), most other papers recognize this cluster as an ICS, and all proposed biosynthetic models and supporting experimental evidence suggest that Dit1 synthesizes an isocyanide intermediate (formyl tyrosine) from l-tyrosine, and thus conforms to our definition of an ICS (13, 93).

Although *dit1* encodes an ICS, this backbone synthase is deeply diverged from all other ICS enzymes (29). Indeed, we confirmed Lim et al. (2018)’s finding that the *dit* ICS is more closely related to bacterial ICSs than to other fungal ICSs (Supplementary Figure S13). This divide is particularly clear when plotting this phylogeny as an unrooted tree, where two distinct clades are evident (Figure 5b). Similar deep phylogenetic divides have been used to infer convergent evolution (94). Although additional research is needed, we speculate that the *dit*-variety fungal ICSs and bacterial ICSs are descendants of a common ancestor, while other fungal ICS enzymes have originated through convergent evolution from a convergently evolved ancestor. Similar patterns of convergence in SM backbone proteins have been documented within plant and microbial terpene synthases (94, 95). To explain the distribution of *dit1* in both fungi and bacteria, we suggest that either an ancient horizontal transfer event occurred or that this cluster is truly ancient and was present in a common ancestor of these two kingdoms. Investigations of bacterial ICS is needed to clarify the ancient evolutionary history of these genes. It is important to note that incomplete lineage sorting or differential selective pressures may have impacted our ability to accurately reconstruct the evolutionary history of ICS enzymes, and thus, the possibility of alternative evolutionary scenarios cannot be entirely ruled out. To shed light on the complicated evolutionary history involving bacteria, a prokaryotic ICS BGC-mining pipeline that does not rely on eukaryotic promoter motifs must be developed.

**Figure 5.**
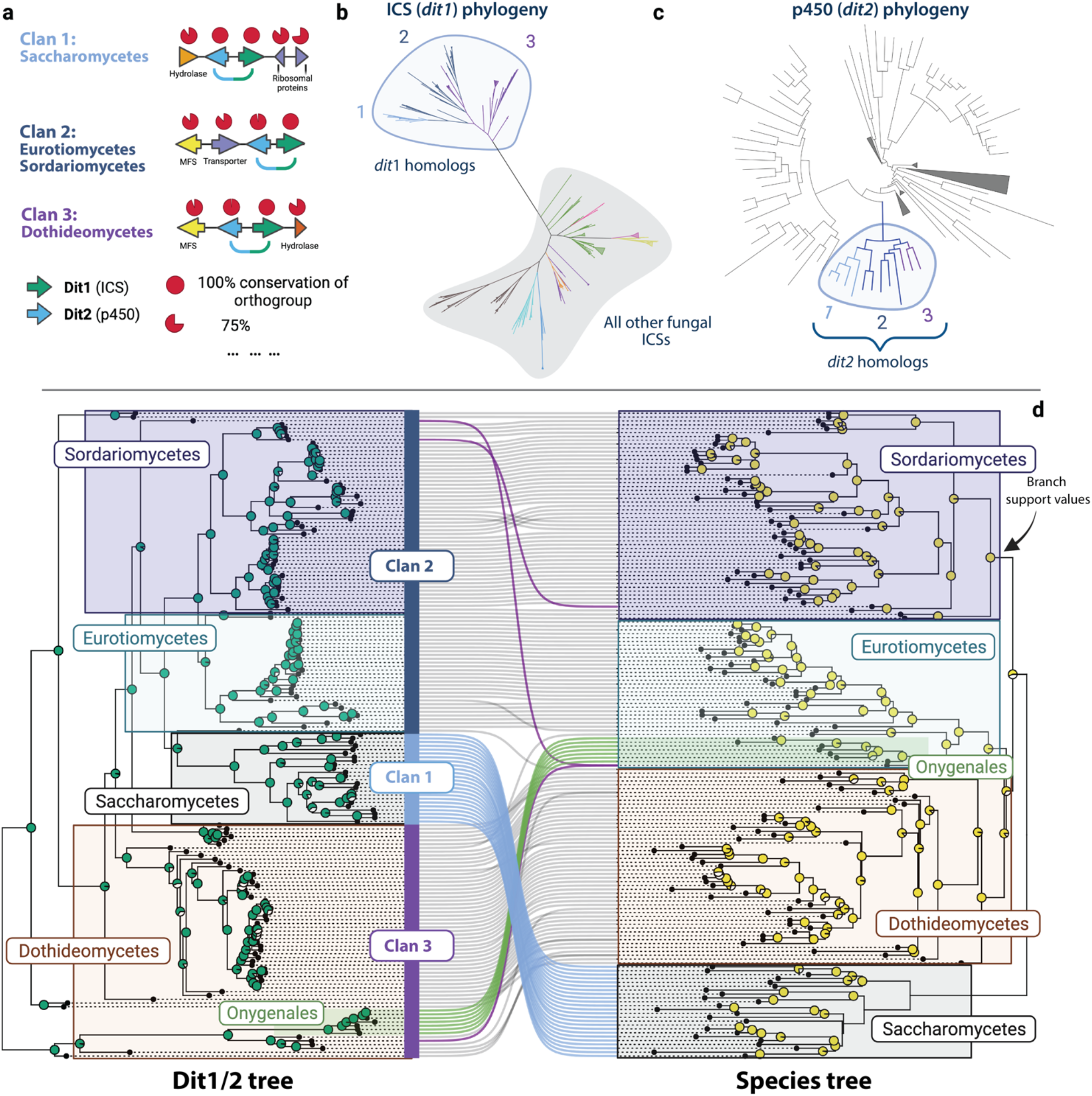
Reconstructed evolutionary history of the *dit* superfamily. (a) The biosynthetic gene cluster (BGC) architectures of the three *dit* clans that make up the *dit* superfamily. The conservation and protein domains for each ortholog is listed above and below the corresponding gene arrow, respectively. The primary phyla each clan is found within is labeled. (b) Unrooted maximum likelihood ICS phylogeny. The ‘lower’ clade’s ICS enzymes (marked in grey) are colored according to the GCF they were found within. All ICSs within the *dit* superfamily (i.e., *dit1* homologs) are found in the ‘upper’ clade (marked with light-blue) and colored according to their clan. (c) The p450 proteins found within the *dit* superfamily BGCs were mapped onto a phylogeny of other p450s found in 10 selected genomes to establish if they are all homologs of *dit2*. As in panel ‘b’, the clans are colored and labeled accordingly. (d) Topological comparison between the concatenated maximum likelihood phylogeny of the Dit1/2 protein sequences (left) and the coalescent-based species tree (right). Branch support values are displayed as pie charts, with the left tree indicating bootstrap support values (1000 bootstrap replicates) and the right tree posterior probabilities. Incongruent phylogenetic relationships are highlighted as follows: (i) Blue: Saccharomycetes, (ii) Green: Onygenales, (iii) Purple: species-level incongruencies. The predominant *dit* clan of each phylum is indicated on the Dit1/2 Tree.

Three primary subclades were identified within the larger *dit* clade of ICS enzymes, all of which were deeply diverged from the ICSs of other GCFs (Figure 5b). The subclades within the *dit* phylogeny largely correspond to three major variations in BGC gene content (denoted as Clans 1-3). The conservation of genes flanking the *dit1*/2 genes could result from genetic hitchhiking, potentially impacting the allele frequencies of neighboring neutral genes (96). However, as flanking genes frequently shared cluster-specific promoters with the *dit1*/*2,* it raises questions about their potential functionality as a coregulated unit. The variation in genomic structure may reflect the divergent evolution of unique functions within specific taxa. Together, the three *dit* Clans constitute what we refer to as the *dit* superfamily (as referred to in Figures 4 and 5). Clan 1 was found primarily in Saccharomycotina, whereas Clan 2 was distributed more broadly across Eurotiomycete and Sordariomycete species, and Clan 3 was predominantly localized in the Dothideomycetes (Figure 5d). These findings shift our understanding of this cluster as it was previously studied in only yeast-like organisms (26, 91, 92). All three Clans are defined by the presence of both an ICS and a p450 whose genes are in the same orientation as the Saccharomycotina *dit1* (ICS) and *dit2* (p450) (Figure 5a). Therefore, we sought to verify that the p450 genes were true *dit2* orthologs as opposed to being independently co-opted p450 proteins (i.e., convergent evolution). Phylogenetic analysis of the putative Dit2 p450s, together with the many other p450s found throughout relevant genomes (including those found in other ICS BGCs), revealed a monophyletic clade of Dit2 sequences distinct from other fungal p450s (Figure 5c). A topological comparison between the Dit1 and Dit2 phylogenies revealed that the two genes closely matched, indicating that the genes have been co-evolving (Supplementary Figure S14). This data suggests that all three *dit* cluster Clans originated from the same ancestral *dit1*/*2* cluster. Additionally, the ubiquitous presence of *dit1* and *dit2* as core genes across all *dit* cluster Clans and taxa provides compelling evidence of their indispensable role in the biosynthesis of *dit* cluster metabolites, regardless of the specific compound(s) produced (i.e., divergent metabolites or conserved dityrosine production).

In total, 17.1% of Ascomycete species in our dataset contained a locus with co-localized *dit1*/*2* homologs. An additional 11.2% of Ascomycetes have a fragmented variation where the *dit1* gene is present in a different genomic locus than *dit2* (Supplementary Table 11). It is uncertain whether the observed fragmentation is entirely due to biological factors or partly due to fragmented genome assemblies. Given the ambiguity surrounding the functionality and divergence of the fragmented *dit* variants, all analyses in this study referencing the *dit* superfamily focused on species that exhibit co-localized loci with *dit1/2*.

### The *dit* superfamily phylogeny is incongruent with the fungal species tree

The *dit* superfamily is the most widely distributed ICS GCF within fungi. We sought to determine the contribution of horizontal and vertical transmission in generating its wide-ranging distribution. Phylogenetic relationships inferred from Dit proteins closely matched those in the species tree among most Sordariomycete, Eurotiomycete, and Dothideomycete species (Figure 5d); this pattern is consistent with vertical transmission and lineage-specific loss. However, all methods used consistently identified two large areas of phylogenic incompatibility. The entire phylum Saccharomycotina, the outgroup to Pezizomycotina in the species tree, was instead the sister clade to the Eurotiomycetes and Sordariomycetes in phylogenies of Dit-cluster proteins (Figure 5d: blue lines). Similarly, Onygenales, an order of dimorphic fungi within the Eurotiomycete in the species tree, was nested within the Dothideomycete clade in the Dit1/2 phylogeny (Figure 5d: green lines). The internal arrangement of species within Saccharomycotina and Onygenales on the Dit phylogeny closely matched the expected species tree arrangement, patterns again consistent with vertical transmission and lineage-specific loss. While we cannot fully rule out the possibility that the short sequences here do not provide full phylogenetic resolution of true evolutionary histories, the incongruency within Saccharomycotina was deeply rooted in the tree and strongly supported by the bootstrap (Dit tree) and posterior probabilities (species tree), while the incongruency within Onygenales was not as deeply rooted but similarly supported by branch support values.

We propose two possibilities to explain the large-scale incompatibilities between the Dit phylogenies and the species tree. First, differences in selection between taxonomic groups could impact phylogenetic relationships through homoplasy (i.e., identity by state, not descent). Unlike most Ascomycetes that have primarily filamentous growth patterns, fungi in some Saccharomyecetes and all Onygenales are considered dimorphic and can exhibit both yeast-like and filamentous growth (97). These lineages are the two largest groups of yeast-like Ascomyecetes. Given that the *dit* cluster is known to be involved in the stability of fungal ascospore cell walls (27, 28, 93), we speculate that the ability of these fungi to grow as yeast could create different selective pressures on cell wall morphology and thus constrain *dit* gene evolution in these species, resulting in phylogenetic incompatibility. The absence of the *dit* cluster in other yeast-like Ascomycetes emphasizes that these genes are not always required for this type of growth. Currently, no Basidiomycete species contained the *dit* cluster. Second, incongruencies in the evolutionary histories of these genes could result from ancient horizontal transfers that would make the evolutionary history of the *dit* genes different from that of the genomes they are found in. Two transfers would have needed to occur, both before the speciation of the Saccharomycotina or Onygenales. Such events would have happened between (i) an ancestor to the Eurotiomycetes/Sordariomycetes and the Saccharomycetes, and (ii) a species in the Dothideomycetes and an ancestor of the Onygenales respectively (Fig. 5d). The Onygenales *dit* cluster shares an MFS gene in the same orientation and order as Clan 3 (Figure 5a), potentially reflecting the history of this putative horizontal transfer.

The *dit1*/*2* cluster in *Saccharomyces cerevisiae* produces dityrosine, an SM incorporated into the protective outer-cell wall layer in its ascospores (26, 27). Deleting *dit2* in *Candida albicans* led to changes in hyphal morphology and increased susceptibility to antifungals, which is believed to be caused by the absence of a dityrosine cell wall layer (91). As reported by Beauvais and Latgé (2018), it remains unclear whether dityrosine is incorporated into the cell walls of filamentous fungi, as current structural models of ascospores are limited to those of *S. cerevisiae*. We found that up to 30% of Ascomycetes possess a co-localized or fragmented copy of *dit1* and *dit2*, with select species (i.e.*, Endocarpon pusillum*, *Botryosphaeria dothidea*) containing multiple copies. Our results raise questions about the potential role of dityrosine in the formation of many Ascomycete ascospores and cell walls. Future studies should aim to characterize the *dit* superfamily SM products across a broader range of taxa, specifically in the Pezizomycotina, to better understand its chemical and ecological significance.

## CONCLUSIONS

Our development of a new genome-mining pipeline has allowed us to identify ICS BGCs across the fungal kingdom, identifying evolutionary patterns that will help inform future research into these compounds. The vast majority of ICSs are part of BGCs that share regulatory motifs (3,800 out of the 4,341). These BGCs are maintained as contiguous clusters over evolutionary time by natural selection, similarly to canonical clusters. ICS clusters are significantly smaller than most canonical clusters, suggesting fewer chemical alterations are made to these compounds. However, the highly-reactive nature of isocyanide intermediates raises the possibility of trans-BGC interactions and yet-unexplored chemistry (14, 20). Indeed, we elucidate that 15.3% of clusters contain synthases from other classes of SMs. While some of these clusters may represent co-localized BGCs that produce chemically distinct products as described for the fumagillin/pseurotin cluster in *A. fumigatus* (89), we speculate that some are true hybrid clusters.

A future goal should be to examine the relationship of isocyanide SMs with metal homeostasis. The reactive nature of isocyanide SMs has been attributed to isocyanide functional group’s ability to interact with metal compounds (19). The *A. fumigatus xan* and *crm* BGCs are regulated by copper and copper transcription factors and their SM products aid in metal-homeostasis and metal-associated competitive interactions (14, 17, 29). We find that the ICS BGCs are three times more numerous than siderophore BGCs, encoding the well-known iron-chelating SMs (99). The unique chemical properties of isocyanide metabolites and the abundance of ICS BGCs identified in this study underscore the potential of these pathways in shaping metal-associated microbial interactions and fungal ecology.

Finally, we find that ICS BGCs are not evenly distributed across fungi, with some taxa, specifically those within the Pezizomycotina, containing disproportionally high numbers of these clusters. While a similar pattern has been observed for canonical gene clusters, we identify that part of this trend can be explained by genome size. Individual GCFs show different distributional patterns, raising questions about the evolutionary histories of these clusters and their role in the niche-adaptation of specific lineages. The most widely distributed ICS GCF was the *dit* superfamily and was found in nearly 30% of all Ascomycetes. The evolutionary history of the *dit* GCF identified deep divergences and phylogenetic incompatibilities that raise questions about convergent evolution and suggests selection or ancient horizontal gene transfers has shaped the evolution of this cluster in some yeast and dimorphic fungi. Our findings highlight important questions for future ICS BGC research aiming to discover and utilize the diverse and valuable ecological properties of isocyanide metabolites (e.g., highly reactive metal-associated microbial natural products) for anthropocentric aims (16, 18, 20). With our web server (www.isocyanides.fungi.wisc.edu), researchers can easily explore, filter, and download all identified ICS BGCs and GCFs, streamlining future research and discovery.

## DATA AVAILABILITY

All the genomes used in this study were obtained from the public domain (NCBI database). Versions of all software used can be found in Supplementary Table S13. The Supplementary Repository contains a reproducible script, every generated ICS BGC prediction, tree files and logs, and additional GCF tables. This can be accessed in the publication’s GitHub repository (https://github.com/gnick18/Fungal_ICSBGCs) or (https://doi.org/10.5281/zenodo.7834839).

## SUPPLEMENTARY DATA

Supplementary Data are available at NAR online.

## ACKNOWLEDGEMENT

We thank Rauf Salamzade for his helpful comments during the creation of this publication. BioRender.com was used in the generation of several figures within this study. Additionally, we thank Nischala Nadig, Dr. Sung Chul Park, and Justin Eagan for their helpful feedback during the development of the webserver. Any opinion, findings, and conclusions or recommendations expressed in this material are those of the authors(s) and do not necessarily reflect the views of the National Science Foundation. Mention of trade names or commercial products in this publication is solely for purpose of providing specific information and does not imply recommendation or endorsement by the US Department of Agriculture. USDA is an equal opportunity provider and employer. Figures 1, 5 and the graphical abstract were created in-part with BioRender.com. The graphical abstract includes some templates generated by BioRender (Citation: *Reprinted from BioRender.com (2023). Retrieved from* https://app.biorender.com/biorender-templates)

## Author Contributions

**Grant R. Nickles**: Conceptualization, Data curation, Formal analysis, Funding acquisition, Investigation, Methodology, Project administration, Software, Validation, Visualization, Writing – Original Draft, Writing –Review & Editing; **Brandon Oestereicher**: Software, Visualization; **Nancy P. Keller:** Conceptualization, Resources, Writing – Review & Editing, Supervision, Funding acquisition; **Milton T. Drott**: Conceptualization, Software, Writing – Review & Editing, Supervision, Project administration, Funding acquisition

## FUNDING

This work was supported by the National Institutes of Health R01 [2R01GM112739-05A1 to N.P.K]; the National Science Foundation Graduate Research Fellowship [Grant No. 2137424 to G.R.N]; the National Institute of General Medical Sciences T32 [GM135066 to G.R.N]; and the United States Department of Agriculture, Agricultural Research Service [to M.T.D]. Funding for open access charge: National Institutes of Health R01 [2R01GM112739-05A1 to N.P.K].

## CONFLICT OF INTEREST

Author N.P.K. declares a potential conflict of interest as co-founder of company Terra Bioforge. N.P.K. is also a Scientific Advisory Board member for Clue Genetics, Inc. The remaining authors declare no competing interests.

## Notes

https://isocyanides.fungi.wisc.edu/

https://github.com/gnick18/Fungal_ICSBGCs

https://doi.org/10.5281/zenodo.7834839

### Competing Interest Statement

Author N.P.K. declares a potential conflict of interest as co-founder of the company Terra Bioforge. N.P.K. is also a Scientific Advisory Board member for Clue Genetics, Inc. The remaining authors declare no competing interests.

