## Supplemental Figures and Methods for "Mining for a New Class of Fungal Natural Products: The Evolution, Diversity, and Distribution of Isocyanide Synthase Biosynthetic Gene Clusters"

**SUPPLEMENTARY FIGURES**

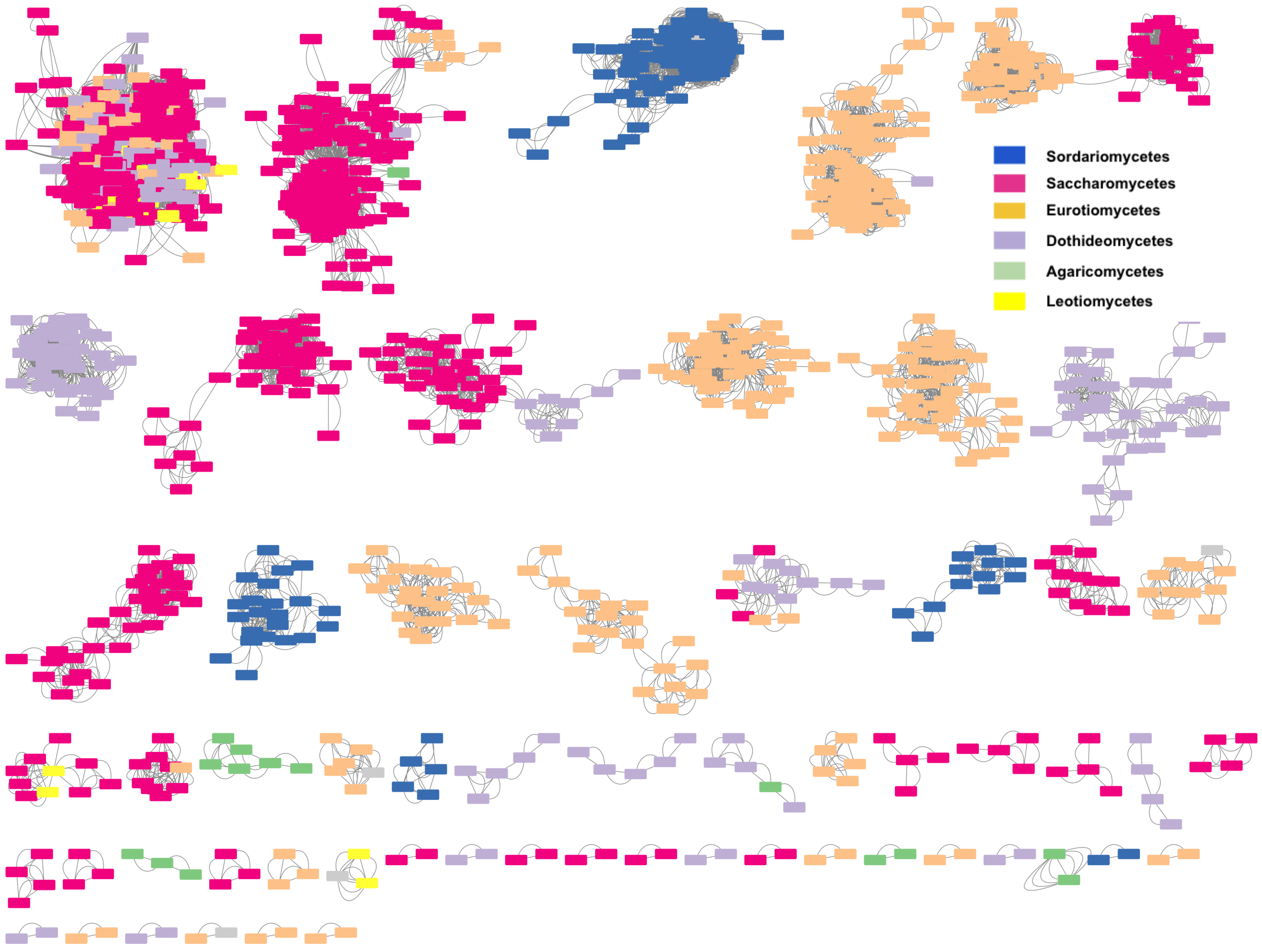


**Figure S1:** Sequence similarity network for a subset of the ICS GCFs. A single BGC prediction is represented as a node and all linked nodes are similar enough to be grouped into a single GCF. Each node is colored based on the taxonomic class it belongs to. Visualization was created with Cytoscape v3.9.1.

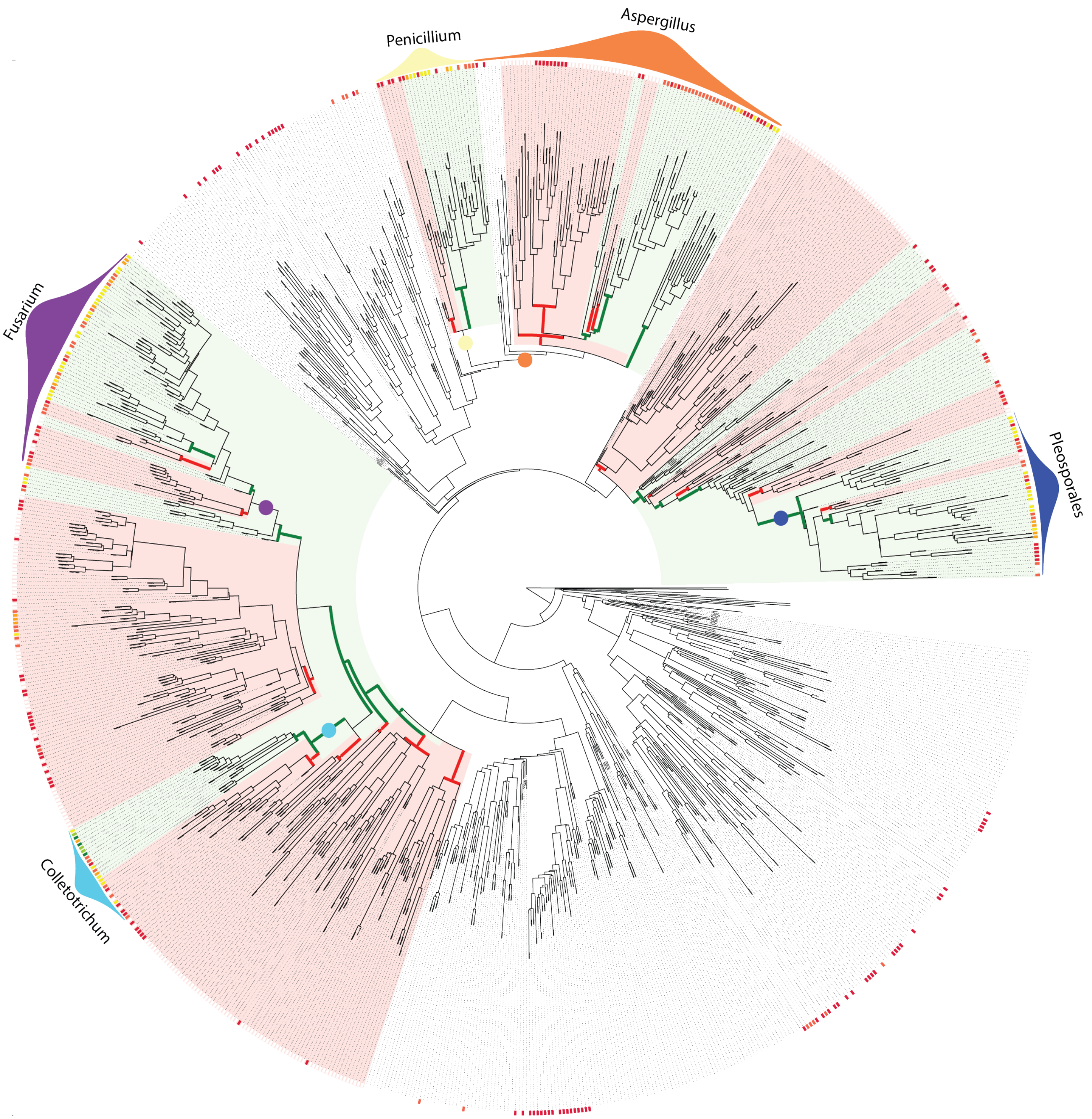


**Figure S2:** Mapping of the potential ICS BGC expansions and contractions within Ascomycota. The outer heatmap on the tree indicates the number of ICS BGCs found in the species, with white indicating none, red 1-2, yellow/orange 3-4, and green >=5. Specific branch points where the sister lineages were found to have statistically significant differences in their mean number of ICS BGCs (T-Tests) are marked as following. (i) Green: The lineage with more ICS BGCs and (ii) Red: The lineage with less ICS BGCs. Certain notable taxa are hand labeled. Please refer to the publication's GitHub repository for an 'svg' version of this tree (every branch is labeled with its species) in addition the raw data from the T-Test results.

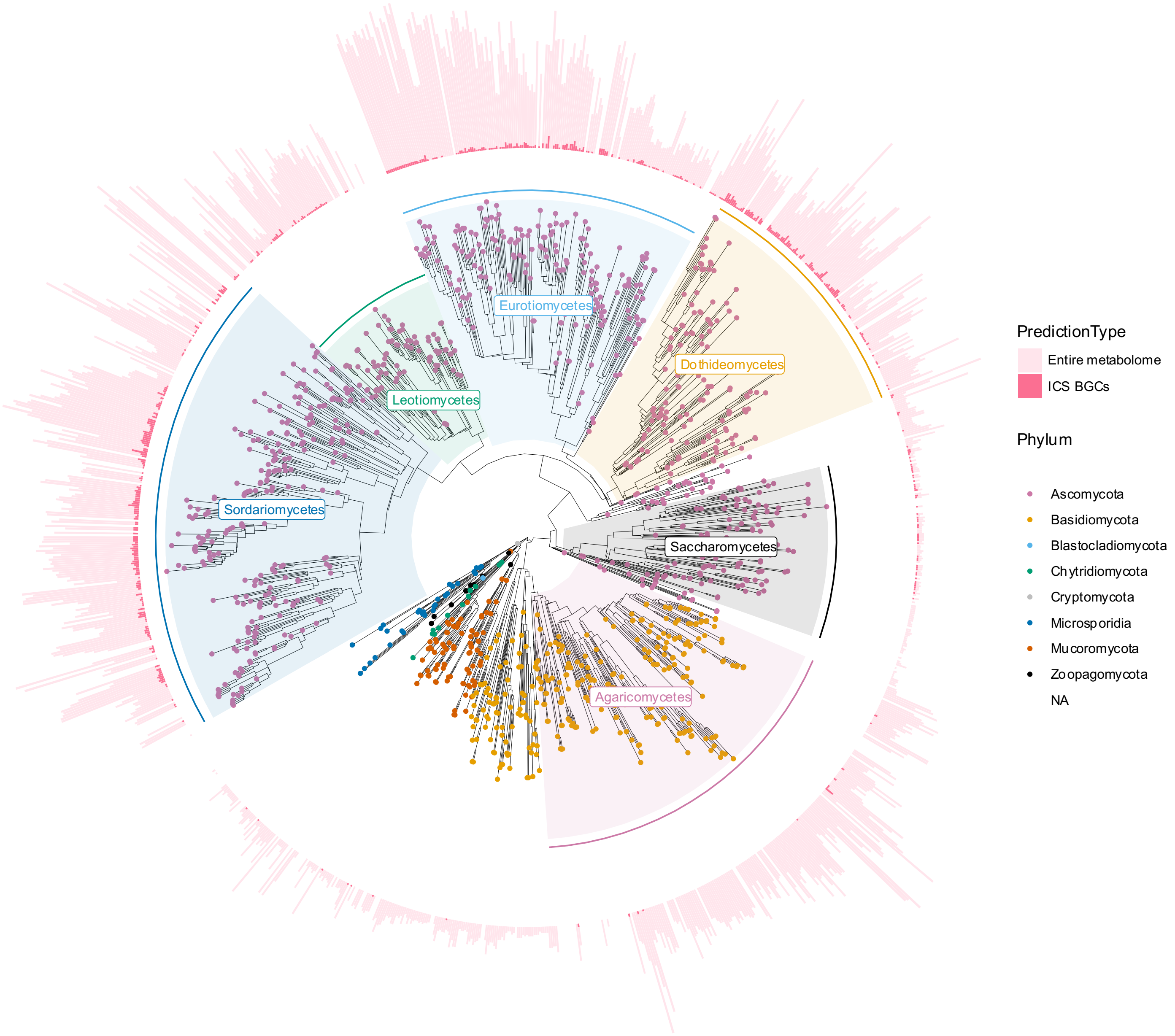


**Figure S3:** The number of ICS BGCs (dark pink), relative to the total predictable biosynthetic potential (light pink) for each species in the dataset.

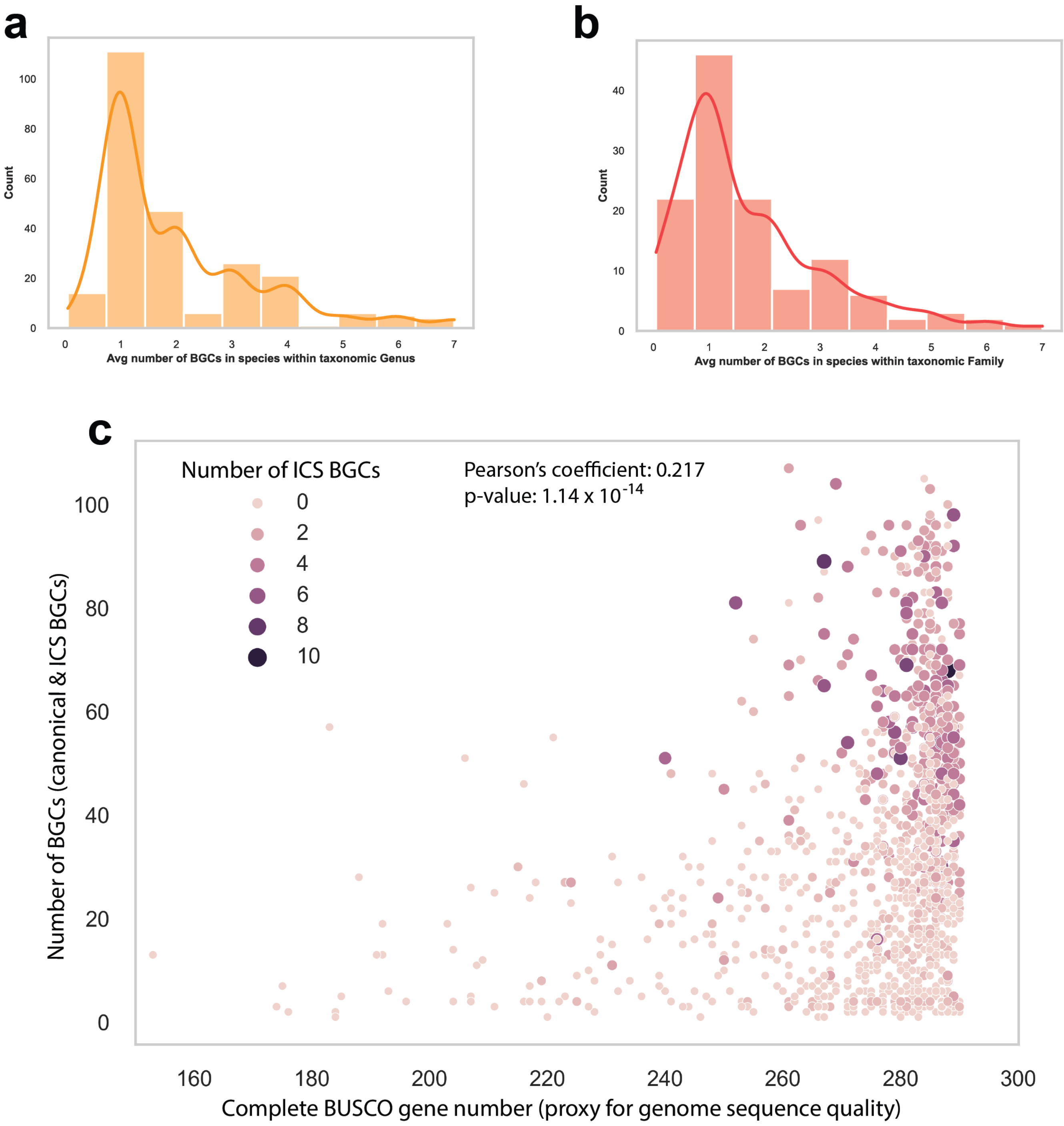


**Figure S4**: Average number of ICS BGCs across taxonomic (a) genera and (b) families. Genera and families with no ICS BGCs were filtered out. Both plots show a positively skewed normal distribution around a peak of 1 ICS BGC. (c) Number of complete BUSCO genes by the number of BGCs (canonical and ICS) within the genomes. The Pearson's coefficient along with the p-value is displayed on the plot.

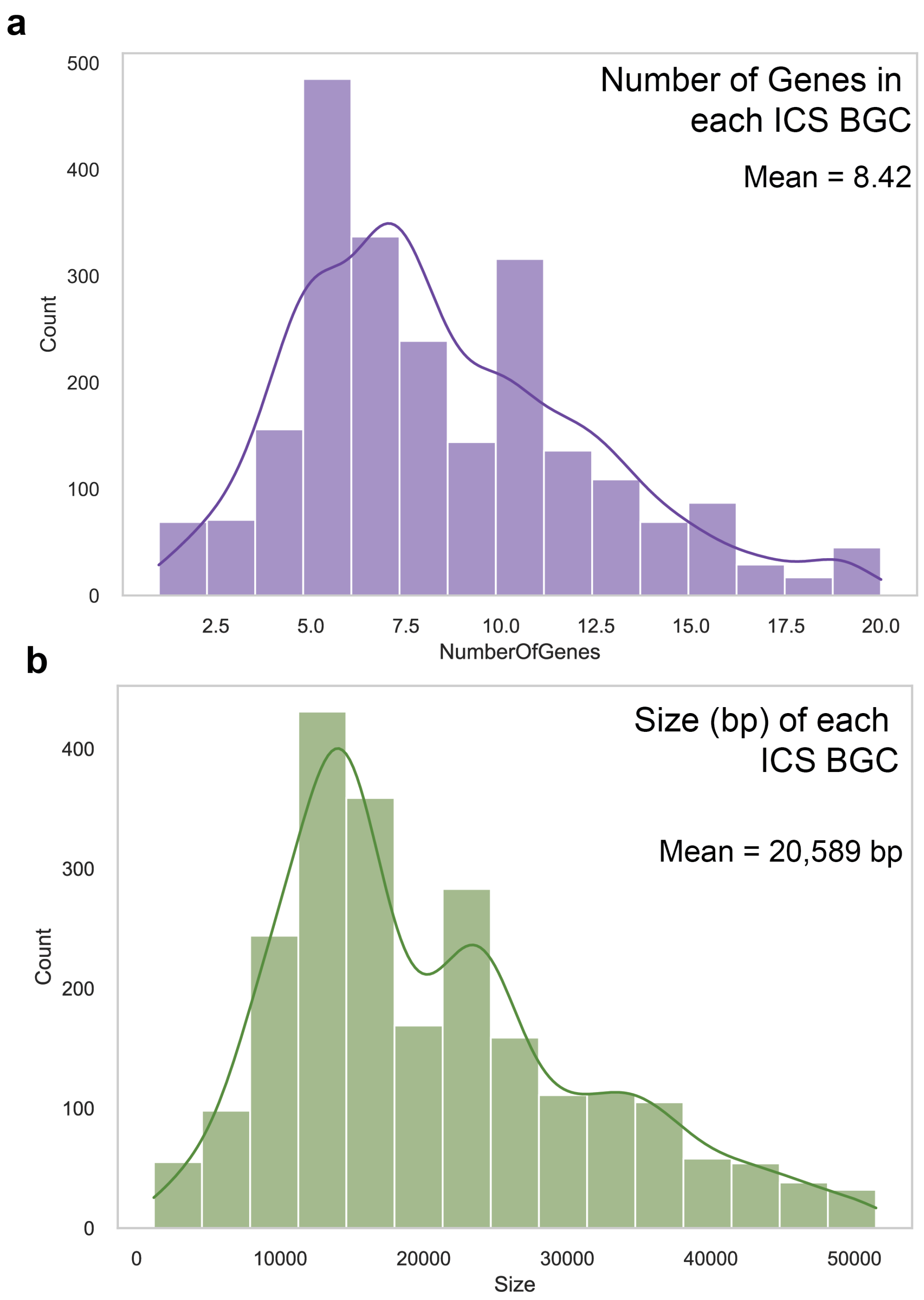


**Figure S5**: The size of the predicted ICS BGCs. Outliers outside of the interquartile range were filtered out. (a) The number of genes is on the x-axis, with count on the y-axis. (b) The size of the entire cluster in base pairs (bp) is on the x-axis, with the count on the y-axis.

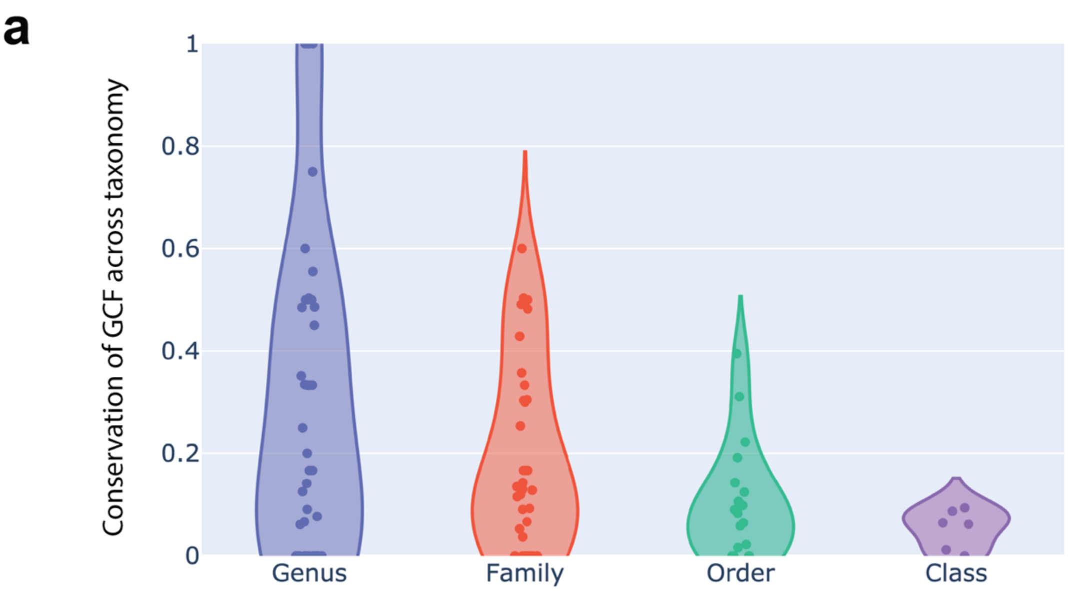


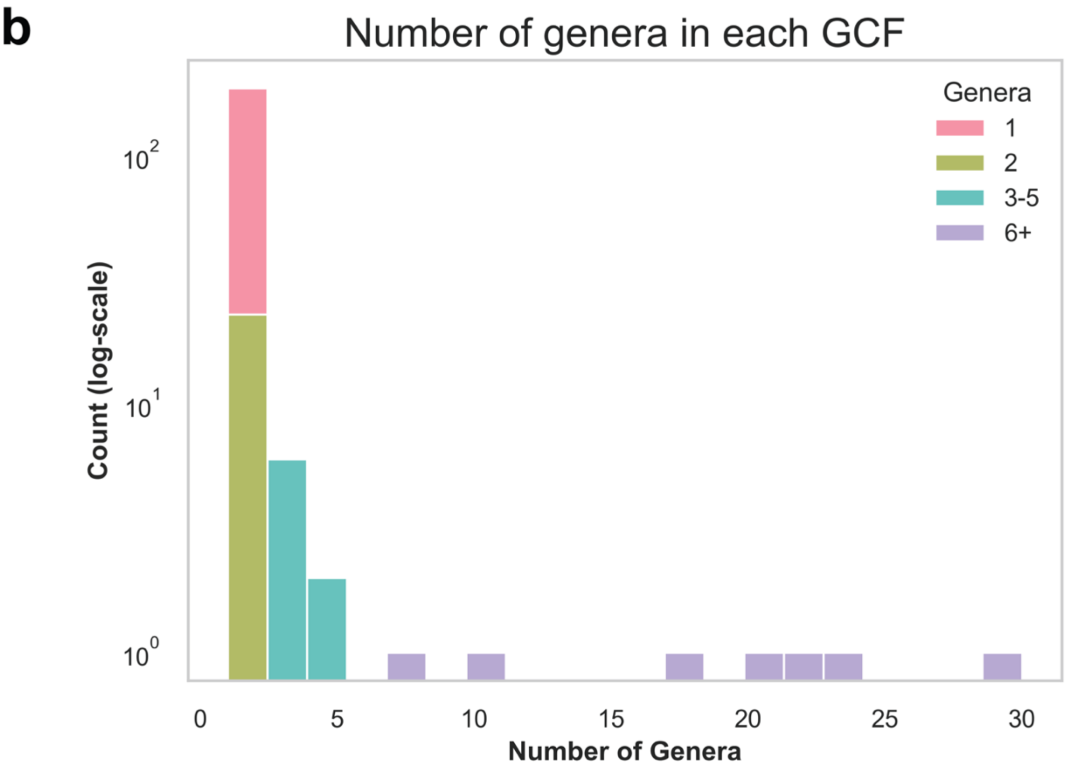


**Figure S6** Distribution of GCFs. (a) Each datapoint indicates the frequency a given GCF was found in species from a given taxonomic rank. GCFs solely found within duplicate isolates of the same species were filtered. Individual datapoints are subject to sampling bias as some of the taxonomies have very poor species coverage. (b) Histogram showing all 200 GCFs and how many genera they were found in. The y-axis is a log-scale of the count, and the x-axis represents the number of genera each GCF was found in. The bars are colored by the number of genera a given GCF contained (see key).

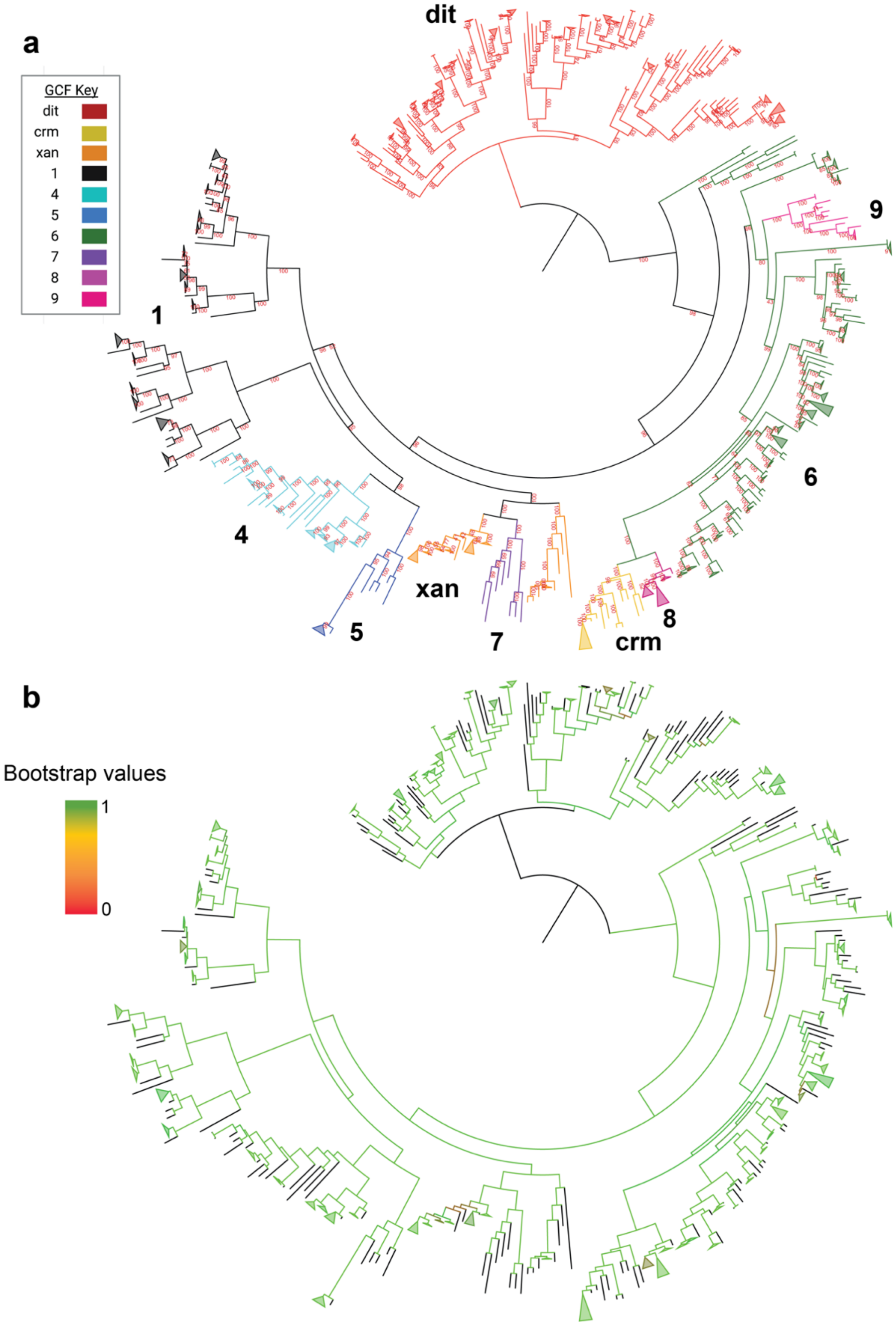


**Figure S7.** Midpoint rooted maximum likelihood phylogeny of ICS proteins from the analyzed GCFs, with bootstrap support values visualized. (a) GCF clades are colored in the same manner as they appear in Figure 4b. Bootstrap values are displayed in red text. (b) The branches are colored with their bootstrap support value using a red-green gradient.

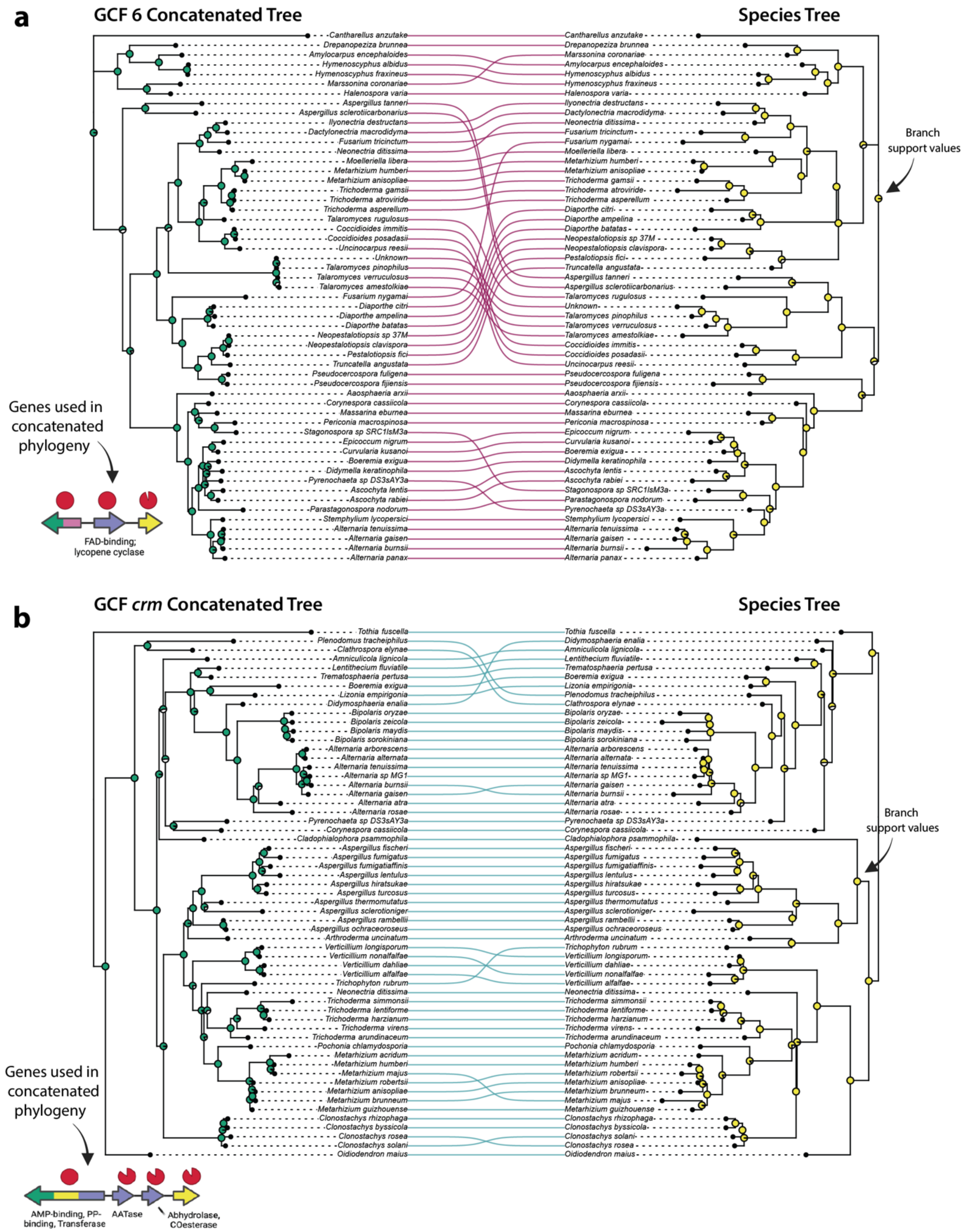


**Figure S8.** Comparisons between GCF 6/crm and the reconstructed species tree. All methods for GCF tree concatenated are identical those described in the making of the *dit* superfamily tree. Only core genes were used in the GCF tree construction, as indicated on each plot. (a) GCF 6. (b) GCF *crm*. Created in-part with [BioRender.com](BioRender.com).

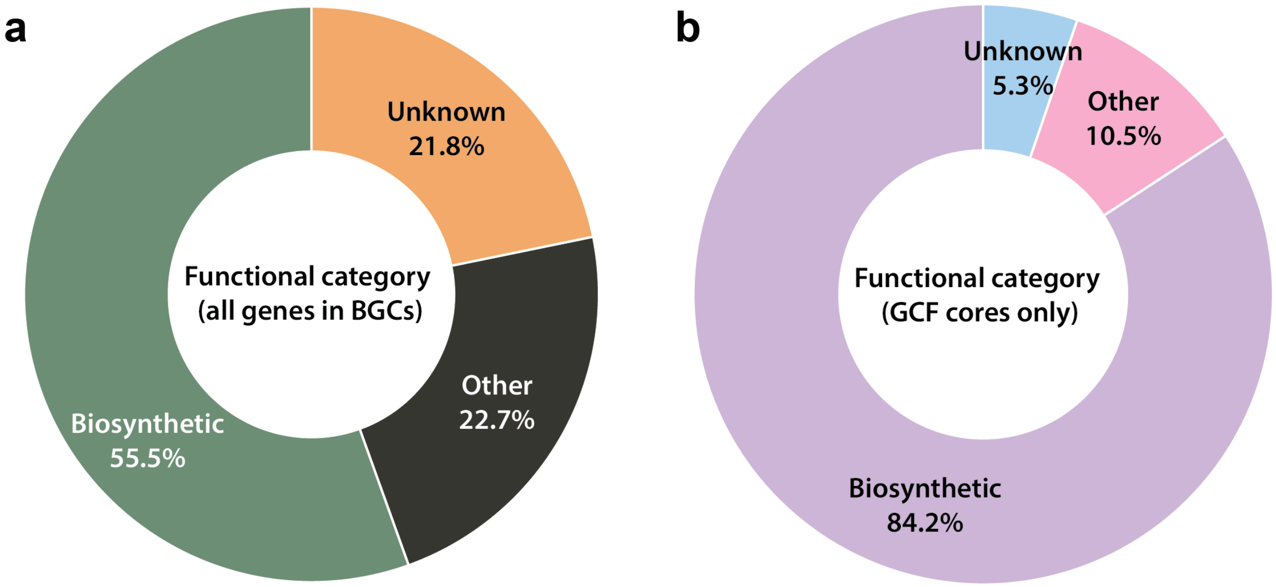


**Figure S9.** Functional categories of the genes found within ICS BGCs. 'Biosynthetic' refers to all genes who contained protein domains prior implicated in SM biosynthesis, transportation, or regulation. 'Other' indicates genes with predictable protein domains not currently known to play a role in SM. 'Unknown' refers to genes with no predictable protein domain. (a) All genes found within the 3800 ICS BGCs. (b) The genes found within the conserved cores of ICS GCFs.

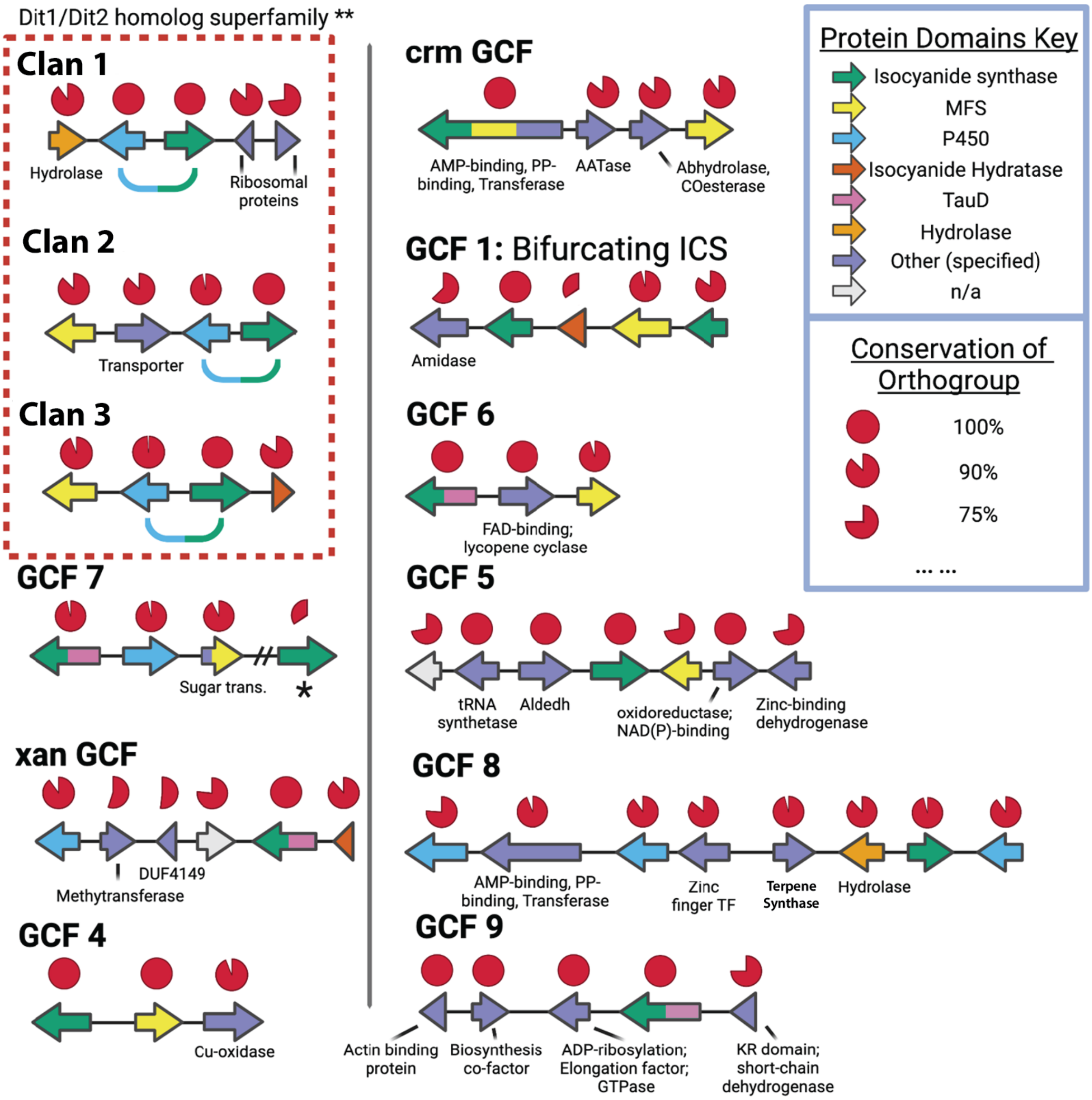


**Figure S10:** The biosynthetic core of each ICS GCF. The conservation of each orthologous gene was calculated by determining the number of species (isolates within the same species were averaged and collapsed) who retained that gene within their copy of said GCF. The values are displayed above each gene as a pie chart. See the key for the color scheme used to denote genes with frequently occurring domains. Genes with GCF specific protein domains are colored in purple and hand-labeled. Created with BioRender.com.

*A second ICS gene was found in around 30% of the GCF 7's BGCs. This second ICS was a homolog of *dit1*.

**The *dit* GCF is characterized by the highly conserved *dit1* (ICS) and *dit2* (p450) genes.

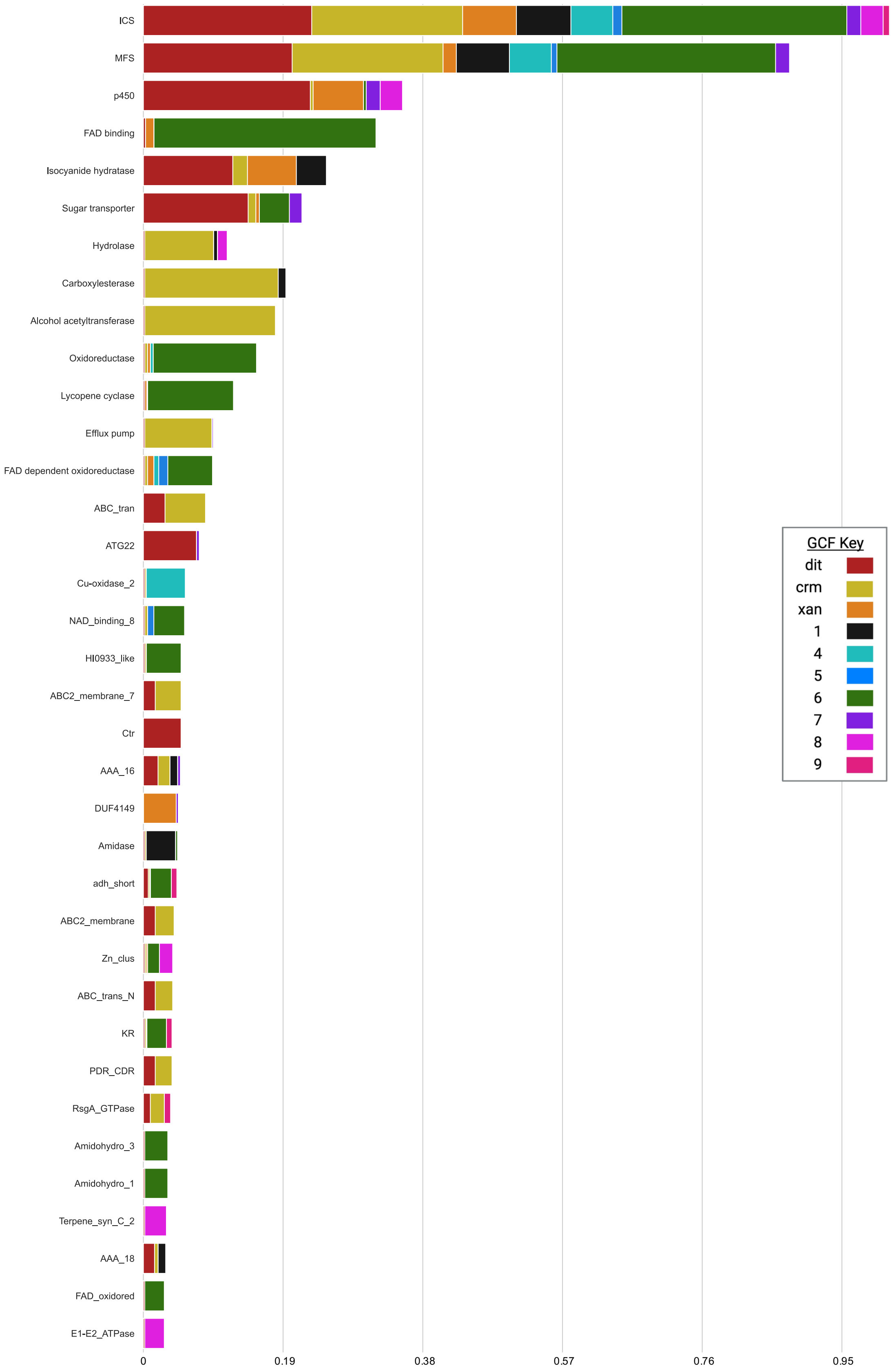


**Figure S11:** The most common protein domains found within the ICS GCFs. This image is a longer version of Figure 4a. The breakdown of different GCFs is colored identically to Figure 4. The names of the domains match the official names in the pFam database.

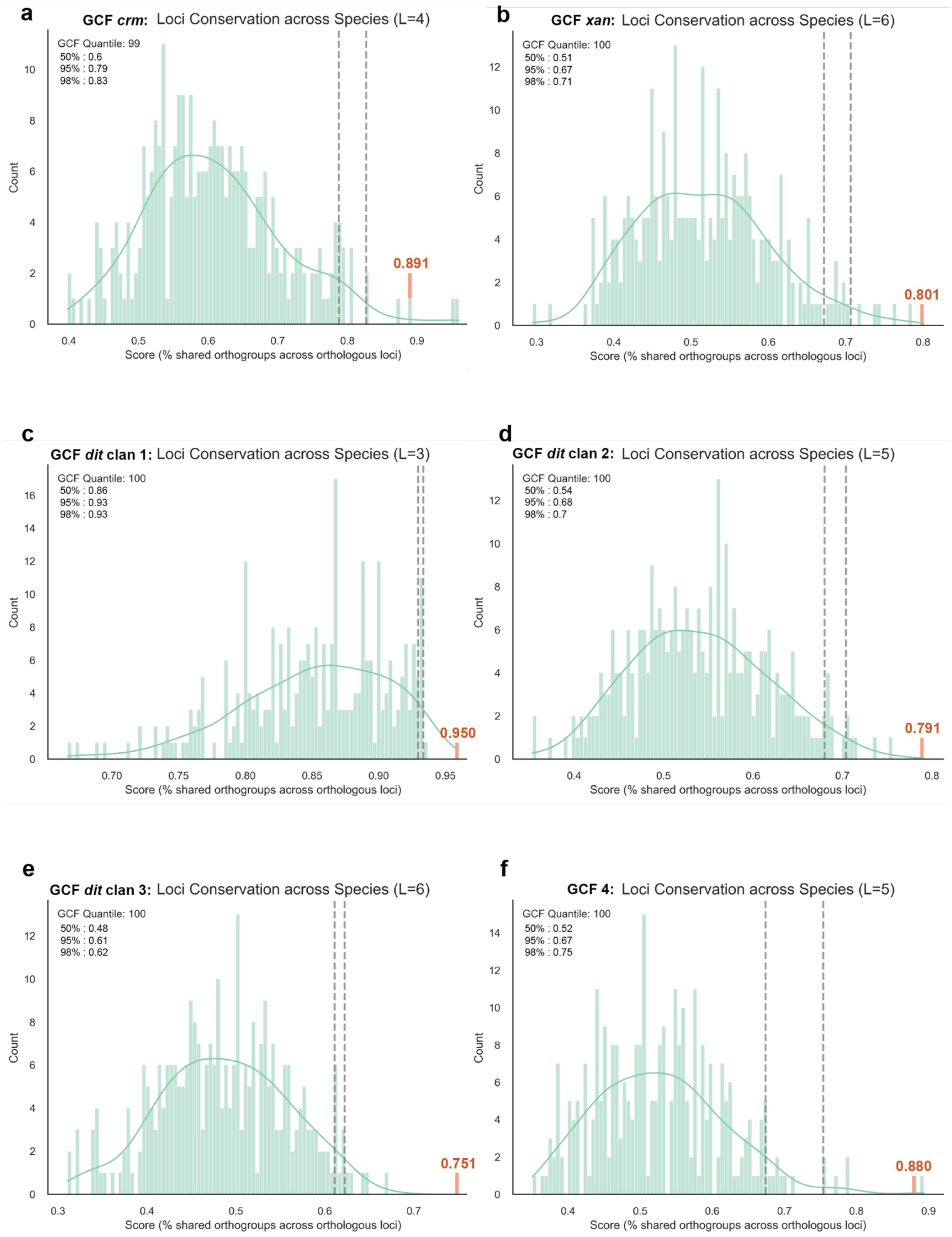
a
GCF crm: Loci Conservation across Species (L=4)
GCF Quantile: 99
50% : 0.6
95% : 0.79
98% : 0.83
0.891
b
GCF xan: Loci Conservation across Species (L=6)
GCF Quantile: 100
50% : 0.51
95% : 0.67
98% : 0.71
0.801
c
GCF dit clan 1: Loci Conservation across Species (L=3)
GCF Quantile: 100
50% : 0.86
95% : 0.93
98% : 0.93
0.950
d
GCF dit clan 2: Loci Conservation across Species (L=5)
GCF Quantile: 100
50% : 0.54
95% : 0.68
98% : 0.7
0.791
e
GCF dit clan 3: Loci Conservation across Species (L=6)
GCF Quantile: 100
50% : 0.48
95% : 0.61
98% : 0.62
0.751
f
GCF 4: Loci Conservation across Species (L=5)
GCF Quantile: 100
50% : 0.52
95% : 0.67
98% : 0.75
0.880
Count
Score (% shared orthogroups across orthologous loci)

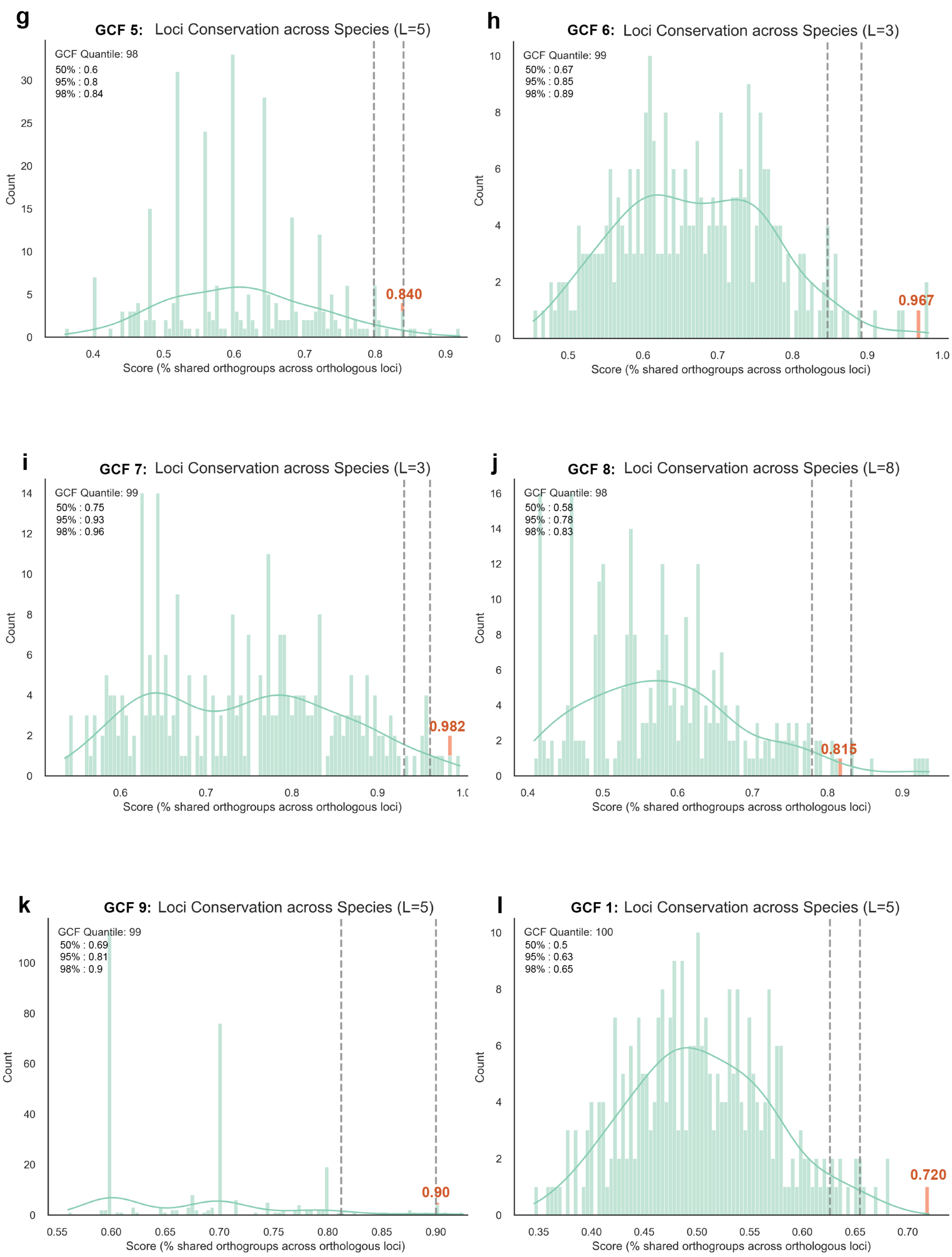
g
GCF 5: Loci Conservation across Species (L=5)
GCF Quantile: 98
50% : 0.6
95% : 0.8
98% : 0.84
0.840
Count
Score (% shared orthogroups across orthologous loci)
h
GCF 6: Loci Conservation across Species (L=3)
GCF Quantile: 99
50% : 0.67
95% : 0.85
98% : 0.89
0.967
Count
Score (% shared orthogroups across orthologous loci)
i
GCF 7: Loci Conservation across Species (L=3)
GCF Quantile: 99
50% : 0.75
95% : 0.93
98% : 0.96
0.982
Count
Score (% shared orthogroups across orthologous loci)
j
GCF 8: Loci Conservation across Species (L=8)
GCF Quantile: 98
50% : 0.58
95% : 0.78
98% : 0.83
0.815
Count
Score (% shared orthogroups across orthologous loci)
k
GCF 9: Loci Conservation across Species (L=5)
GCF Quantile: 99
50% : 0.69
95% : 0.81
98% : 0.9
0.90
Count
Score (% shared orthogroups across orthologous loci)
l
GCF 1: Loci Conservation across Species (L=5)
GCF Quantile: 100
50% : 0.5
95% : 0.63
98% : 0.65
0.720
Count
Score (% shared orthogroups across orthologous loci)

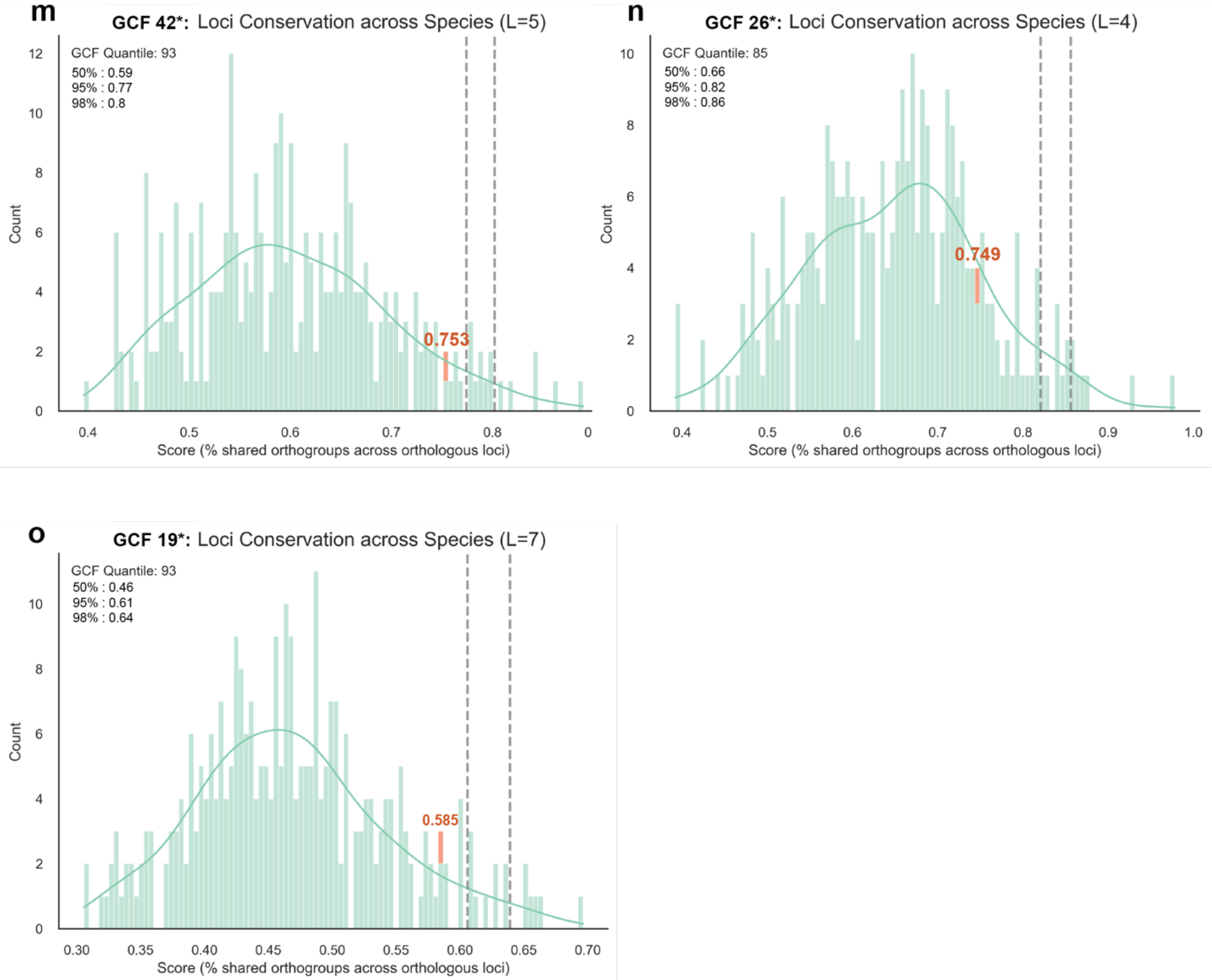


**Figure S12:** Estimating the extent of linkage disequilibrium (LD) within the 15 tested GCFs by comparing the conservation of syntenic genes around BUSCO genes and GCF loci. The conservation scores (e.g., percentage of shared orthologs in a given locus across species) for the whole-genome rates of gene drift are shown in green. 'L' refers to the number of genes used in the individual GCF's analysis and was determined by the size of the GCF's biosynthetic core. The conservation score for each GCF locus is highlighted in orange. The two dashed grey lines indicate the threshold for the 95$^{th}$ and 98$^{th}$ percentiles, which are also listed in the top left. Panels L-N are GCFs tested that did not have values above the 95$^{th}$ percentile, and still have their original GCF names from the BiG-SCAPE dereplication pipeline.

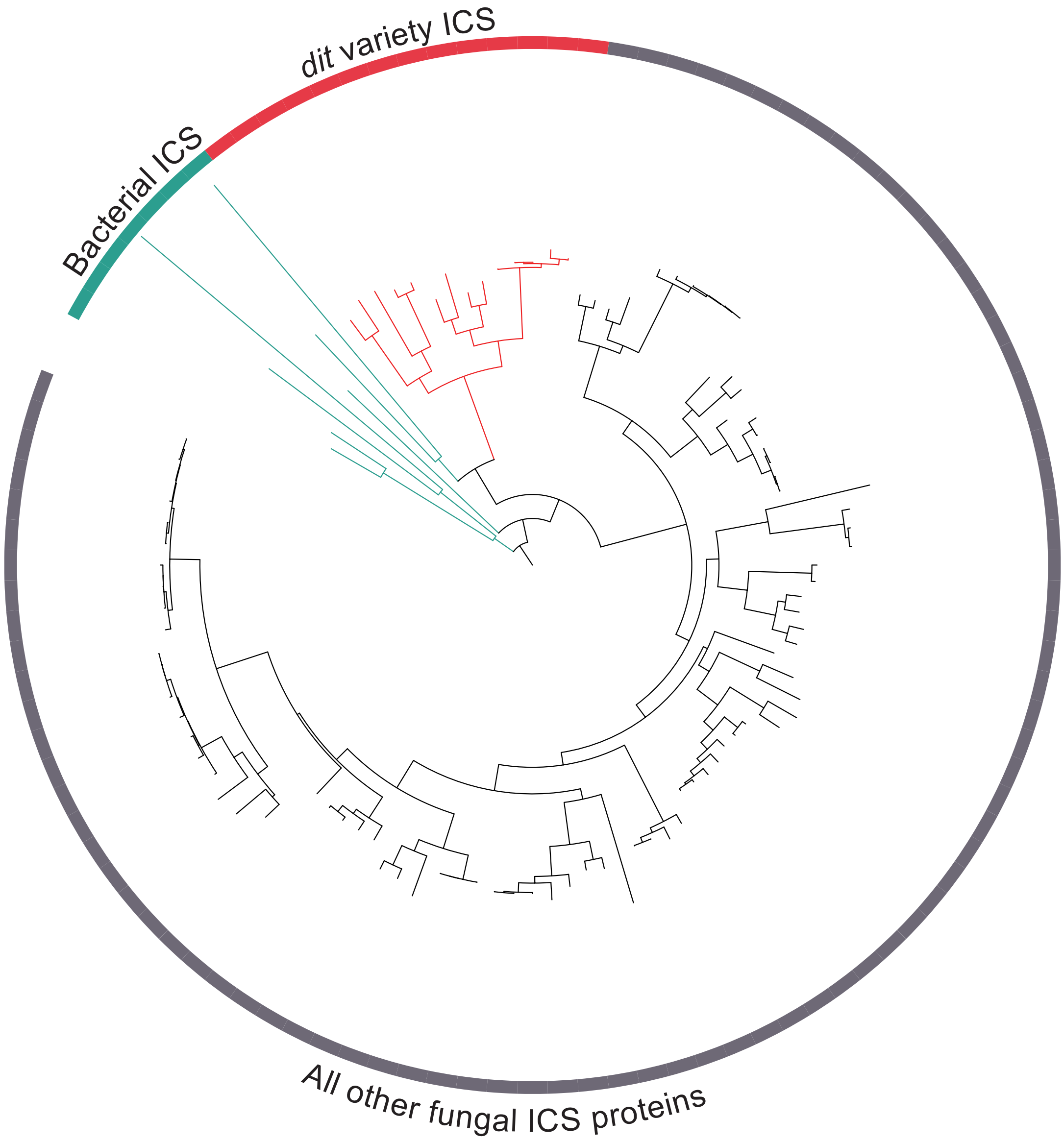


**Figure S13**: The relatedness of fungal ICS proteins to a sampling of bacterial ICS. The midpoint-rooted tree was constructed using a representative sampling of fungal and bacterial ICS genes.

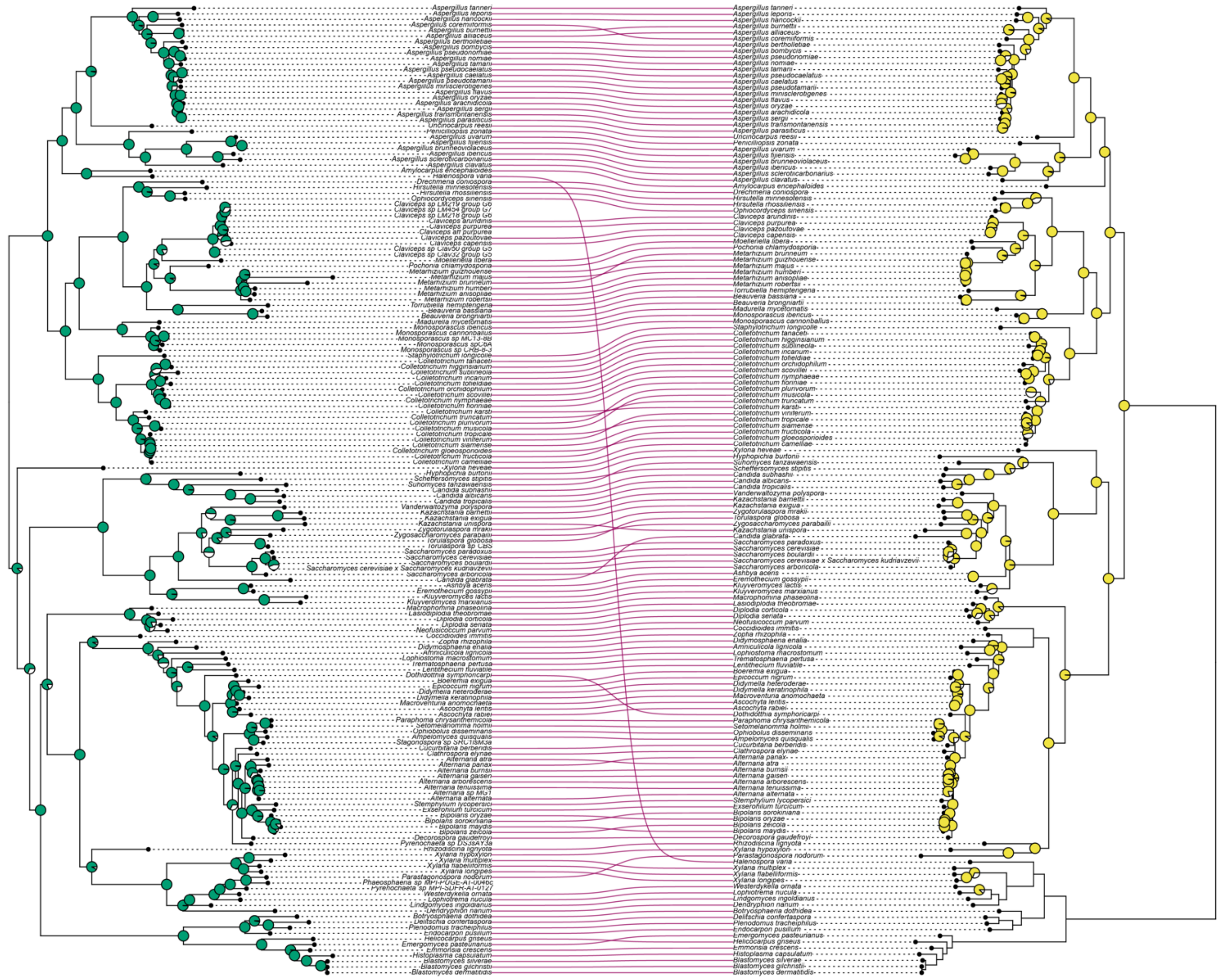

**Figure S14.** Topology comparison between the Dit1(ICS) and Dit2 (p450) phylogeny to identify patterns of co-evolution.

**SUPPLEMENTARY METHODS:**

**Calculating the proportion of the fungal biosynthetic potential that ICS BGCs account for**

After running antiSMASH v5 (1) on all 3,300 genomes, the backbone enzyme defined BGC classes were sorted in accordance with the antiSMASH defined classifications. The decision was made to use only one representative genome for each species to prevent species who have numerous sequenced isolate genomes from skewing the overall results. The isolate with the largest number of predicted canonical BGCs for any given species was calculated to make this determination. While we do acknowledge that this choice could bias the results for highly fragmented genomes with multiple BGC predicted in split contig BGCs, this would have inconsequential impacts on the overall analysis being conducted We verified this by calculating the percent of the metabolome that ICS BGCs make up in the unfiltered and filtered datasets. The overall values for each BGC group fluctuated by at most 2 percentage points, and the order of the group stayed consistent throughout. As an example, ICS BGCs make up 4.49% of the specialized metabolome in the unfiltered dataset, and 3.0% in the filtered dataset.

**Dereplicating the GCF output of BiG-SCAPE**

The accuracy of GCF groupings is dependent on the accuracy of BGC border predictions (i.e., what neighboring genes are included). Miss-calls of BGC borders can lead to incorrect GCF classifications. Part of this issue arises when distinct GCFs are linked together by a single or small number of BGCs (that we dub 'bridge clusters' as these calls often connect two distinct GCFs), leading to their inappropriate classification as a single GCF (i.e., a single BGC linking two separate GCFs). Bridge clusters are often explained by the miss-prediction of orthologous neighboring genes shared between otherwise distinct BGC predictions. The output of BiG-SCAPE underwent systematic dereplication resulting in the separation of bridge clusters into individual GCFs ('Step 5' in Reproducible Script). Refer to Supplementary Table S5 for details on the dereplicated GCF categorizations for each ICS BGC prediction.

**Identifying putative expansions and contractions**

The species tree created in this publication (Fig. 2) was subset to only contain Ascomycota species and a single Basidiomycete outgroup (*Amanita muscaria*). A custom Python program was developed to identify potential expansions and contractions in the number of ICS BGCs between sister clades using independent t-tests (2). The program is designed to start its search from a node of choice provided by the user. Every branch point beneath said node is then explored. For each branch, the program extracts the number of ICS BGCs in the sister clades and conducts a t-test to detect significant differences below the user-defined threshold e-value. The program then generates an output file containing the t-test results, and produces visualizations of the expansions and contractions as can be seen in Fig. S6. The script to run this Python pipeline is included in the publications Reproducible Scripts file, and the data from the t-tests on ICS BGCs along with the tree are available on the publication's GitHub repository.

While this type of analysis has its merits, it is important to note its limitations. One major concern is that statistical tests comparing the equivalence of two groups does not take into account the time elapsed since the divergence of two taxa, which can result in misinterpretation of results due to gene drift rather than selection (3). To address this issue, stochastic birth and death models have been proposed, which compare selection to the null model of random gene birth and loss (4, 5). However, accurate time-dated species trees are required for such models, which are difficult to construct for fungi due to low number of fossil records and high-quality genomic sequences (6, 7). Such programs also assume the most recent common ancestor contains a single copy of the studied gene or cluster of genes. It remains uncertain whether assuming the most recent common ancestor of all Ascomycetes contained an ICS BGC is a valid assumption given the evidence of convergent evolution between two separate clades of ICS backbone synthases (as depicted in Fig. 5). It was because of these limitations that we developed our own program using traditional statistical tests. While our approach does provide a crude identification of key points on the tree where large differences exist, further research is needed to determine the cause of such putative expansions or contractions.

**Kendall’s tau correlation analysis between the number of ICS BGCs and canonical BGCs**

SciPy v1.5.2 (8), researchpy (9), ETE 3 v3.1.2 (10), and pandas v1.3.4 (11) were all used for this analysis. We performed a Kendall's tau correlation analysis (12) to examine the relationship between the number of ICS BGCs identified in fungi and the corresponding number of canonical clusters. The SciPy v1.5.2 (8) command ‘kendalltau’ was used for this analysis with the default ‘tau-b’ variant.

**Determining what proteins were biosynthetic, unknown, or other**

We adopted the same criteria for defining "biosynthetic" protein domains as the widely-used BGC predictive software antiSMASH v5 (1). This included all enzymatic, regulatory, and transportation domains (e.g. MFSs) that have been previously reported to function within BGCs. Using such criteria, we categorized each gene into one of three groups: (i) "Biosynthetic" if it contained at least one biosynthetic domain, (ii) "Other" if it contained protein domains not previously associated with SM biosynthesis, or (iii) "Unknown" if no predictable protein domain was detected.

**Identifying homologous sets of co-localized genes using cBlaster**

A representative BGC prediction was hand-selected for each of the 31 analyzed GCFs. This selection was based on a BGC containing all the highly conserved orthologs for a given GCF. To make more informed choices for the selection of a representative BGC for each GCF, we used Orthofinder v2.5.2 (13) to identify the shared set of orthologous genes within each GCF. This enabled us to locate which orthogroups were highly conserved across all the BGCs within a GCF. The representative BGC for each GCF was chosen if it contained all the highly conserved orthologs for said GCF.

We utilized the rapid identification of homologous gene clusters tool cblaster v1.3.15 (14) to search for homologous loci to the query sequence. A local database from genomic GenBank files was created using the 'makedb' module, which implements the alignment software DIAMOND (15). The database was filtered down to include a maximum of 3 isolates for any given species to increase run speeds. This selection was done at random. The following pipeline was then employed for all 31 GCFs: (ii) The session for the default run was used to create gene neighborhood estimations using the 'gne' module, allowing for the optimal intergenic distance value to be calculated. (iii) An optimized search was re-run, using the calculated intergenic distance threshold, a list of backbone ICS genes the hit must include, and the minimum number of hits a species should have for the result to return the putative cluster (was adjusted for unique size of each GCF). Overall, this second approach to identifying BGCs is independent of regulatory elements and allowed us to fine-tune our CASSIS predictions around the highly conserved GCF cores.

**Refining the list of species with the *dit* cluster**

We used a phylogenomic approach to verifying if species had a homolog to the dit cluster. Briefly, all ICSs that fell within the *dit* clade on the ICS protein phylogeny were extracted. If such gene was not originally predicted to be within an ICS BGC using the CASSIS algorithm, we searched to check if a copy of *dit2* was within 2 genes on the *dit1* homolog. In cases where the *dit2* gene was fragmented from *dit1*, it was documented but not included in any subsequent analyses. The results of the verification step, in addition to the protein names of any hand identified *dit* clusters can be found in Supplementary Table S11.
